# Giant viruses specific to deep oceans show persistent presence and activity

**DOI:** 10.1101/2025.06.21.660676

**Authors:** Wenwen Liu, Komei Nagasaka, Junyi Wu, Hiroki Ban, Ethan Mimick, Lingjie Meng, Russell Y. Neches, Mohammad Moniruzzaman, Takashi Yoshida, Yosuke Nishimura, Hisashi Endo, Yusuke Okazaki, Hiroyuki Ogata

**Author notes:** Corresponding author: Hiroyuki Ogata,.

## Abstract

Giant viruses (GVs) of the phyla *Nucleocytoviricota* and *Mirusviricota* are large double-stranded DNA viruses that infect diverse eukaryotic hosts and impact biogeochemical cycles. Their diversity and ecological roles have been well studied in the photic layer of the ocean, but less is known about their activity, population dynamics, and adaptive strategies in the aphotic layers. Here, we conducted eight seasonal time-series samplings of the surface and mesopelagic layers at a coastal site in Muroto, Japan, and integrated 18S metabarcoding, metagenomic, and metatranscriptomic data to investigate deep-sea GVs and their potential hosts. The analysis identified 48 GV genomes including six that were exclusively detected in the mesopelagic layer. Notably, these mesopelagic-specific GVs showed persistent activity across seasons. To investigate the global deep-sea-specific GV distribution, we compiled GV reference genomic data from the OceanDNA MAG project and other resources, and analyzed 1,890 marine metagenomes. This revealed 101 deep-sea-specific GVs, distributed across the GV phylogenetic tree, indicating that adaptation to deep-sea environments has occurred in multiple lineages. One clade enriched with deep-sea-specific GVs included one GV identified in our Muroto sampling, which displayed a wide geographic distribution. Seventy-six KEGG orthologs and 74 Pfam domains were specifically enriched in deep-sea-specific GVs, encompassing functions related to the ubiquitin system, energy metabolism, and nitrogen acquisition. These findings support the scenario that distinct GV lineages have adapted to hosts in aphotic marine environments by altering their gene repertoire to thrive in this unique habitat.

**IMPORTANCE:** Giant viruses are widespread in the ocean surface and are key in shaping marine ecosystems by infecting phytoplankton and other protists. However, little is known about their activity and adaptive strategies in deep-sea environments. In this study, we performed metagenomic and metatranscriptomic analyses of seawater samples collected from a coastal site in Japan and discovered giant virus genomes showing persistent transcriptional activity across seasons in the deep-sea water. Using a global marine dataset, we further uncovered the widespread existence of deep-sea-specific giant viruses and characterized their unique gene repertoire, which likely facilitates adaptation to the limited availability of light and organic compounds in the aphotic zone. These findings expand our understanding of giant virus ecology in the dark ocean.

## INTRODUCTION

Giant viruses (GVs), encompassing the phylum *Nucleocytoviricota* and the recently proposed phylum *Mirusviricota*, are characterized by their large virion and genome sizes (1–3). They infect a broad range of eukaryotes from unicellular algae, diverse heterotrophic protists, to larger animals (4–6). In marine environments, GVs affect the population dynamics of their hosts including bloom-forming algae (7–9), alter host metabolism via virus-induced metabolic reprogramming (10, 11), and consequently impact carbon and nutrient cycles (12). Previous metagenomic studies have revealed the high abundance (13), activity (14, 15), diversity (16–18), and widespread geographic GV distribution in the ocean (19–21). The community structure of GVs in marine environments tightly correlates with that of microeukaryotes in time (22–24) and space (25), supporting the idea that GVs regulate the microeukaryote community.

To date, studies have predominantly focused on the ocean’s photic layer, while aphotic layers—which constitute approximately 95% of the global ocean—remain comparatively underexplored owing to the inherent difficulty in accessing these environments. Of note, all marine GVs have been isolated from the photic layer, except for one case from deep-sea sediment (26). Several previous studies nevertheless detected the presence of GV-related sequences in deep-sea environments, including genomic contigs (27–29) and conserved marker genes (19, 30). Another study reported the transcription signals from mirusvirus genomes in mesopelagic waters based on *Tara* Oceans data (2). More recently, five GV metagenome assembled genomes (GVMAGs) were shown to be unique to the deep water below 150 m (31) and a few GVMAGs were specifically and highly abundant in the deep layers of the North Pacific Subtropical Gyre (29). These results clearly indicate the existence of GVs in deep oceans. However, whether these GVs represent deep-sea residents actively infecting sympatric hosts or are just passively transported from the surface community through sinking remains unclear. For example, Endo *et al*. showed that the GV lineages detected in the mesopelagic layer are mostly (99%) also present in the photic layer and suggested that they are largely settled from the photic layer (19). Sheam *et al*. also provided evidence that some GVs detected in sediment trap samples (4,000-m depth) were vertically transported from the surface layer (29). Nonetheless, a recent study revealed the existence of a GV community specific to an aphotic layer (65-m depth) of a freshwater lake (32), implying the existence of marine GVs specifically inhabiting dark environments.

Investigating GVs in aphotic layers is challenging because of methodological and logistical difficulties (33), as GVs have lower abundance than bacteria in the same size fraction, especially in deeper layers (13). Thus, a large volume of water must be sampled to concentrate the GVs. Furthermore, collecting samples for transcriptomic analysis to probe viral activity needs to be conducted in a relatively short time to avoid RNA degradation and shifts in the physiological state of the microorganisms (34). The Kochi Prefectural Deep Seawater Laboratory, located in Muroto, Kochi Prefecture, Japan, is situated at a coastal site, where oligotrophic water influenced by the Kuroshio Current can be readily accessed (35). The laboratory has the facility to continuously draw untreated surface (0.5 m) and mesopelagic (320 m) seawater (36), and supports scientific research of deep-sea water and various industrial applications such as food processing and aquaculture (37).

In this study, we applied metabarcoding, metagenomic, and metatranscriptomic analyses of the surface and mesopelagic samples collected across seasons at the Laboratory in Muroto to investigate the existence and activity of GVs, as well as the sympatric potential host community, in the mesopelagic layer. We identified and characterized 48 GVMAGs, 6 showing stable activity exclusively in the mesopelagic water. We then compiled a global GVMAG dataset, named the Giant Virus Genome Reference (GVGR) database, from publicly available data (2, 6, 23, 32, 38, 39), and performed a systematic analysis to identify deep-sea-specific GVs at a global scale. The analysis revealed 101 GVMAGs preferentially detected in deep oceans, some forming deep-sea-specific GV clades in the phylogenetic tree.

## RESULTS AND DISCUSSION

### Data overview

To investigate the communities of GVs and microeukaryotes in the surface and mesopelagic layers at the study site (i.e., off the coast of Muroto), we generated 18S rDNA and rRNA metabarcoding (4,981 ASVs in total), metagenomic (757 Gbp in total), and metatrancriptomic (598 Gbp in total) data from seawater samples collected from two depths (0.5 m and 320 m) (Fig. S1, Table S1|). The pump systems at the Muroto Laboratory (Fig. S1b) enabled us to collect and filter a large volume of water in a short time (Table S2). For metabarcoding and metatranscriptome analyses, we used samples of two size fractions: the “small” size fraction (0.2–3.0 μm, 0.2–5.0 μm, or 0.2–150 μm) and the “large” size fraction (3.0–150 μm or 5.0–150 μm). For metagenomic analysis, we used the samples from the small size fraction. From these samples, we successfully extracted adequate quantities (> 500 ng) of high-quality DNA and RNA for sequencing (Table S3).

### Mesopelagic microeukaryote community is distinct from that in the surface water at the sampling site

In this study, 18S rDNA metabarcodes at the ASV level were used to measure approximate organism abundances, although the abundance of 18S rDNA is biased by the copy number of rRNA genes, which varies among organisms (40, 41). 18S rRNA metabarcodes were used as a proxy for the metabolic activity of organisms (42). The microeukaryote community in the mesopelagic layer substantially differed from that of the surface layer at the study site (Fig. 1a–d), suggesting that the two layers harbor distinct host candidates for GVs. Furthermore, the mesopelagic microeukaryote community characterized by 18S rRNA differed from that of 18S rDNA metabarcodes, suggesting that some microeukaryotes are active in the mesopelagic layer, while others are inactive (i.e., in a dormant state or dead cells settled from the surface). Non-metric multidimensional scaling (NMDS) ordination confirmed a clear separation between the surface and mesopelagic microeukaryote communities, regardless of whether 18S rDNA (Fig. 1e; R^2^ = 0.25, PERMANOVA, *P* < 0.01) or 18S rRNA (Fig. 1f; R^2^ = 0.24, PERMANOVA, *P* < 0.01) metabarcodes were used. It was noted that the mesopelagic microeukaryote community showed a lower level of seasonal variation, providing a more stable long-term host environment for GVs compared with the surface community (Fig. 1e,f).

**Figure 1.**
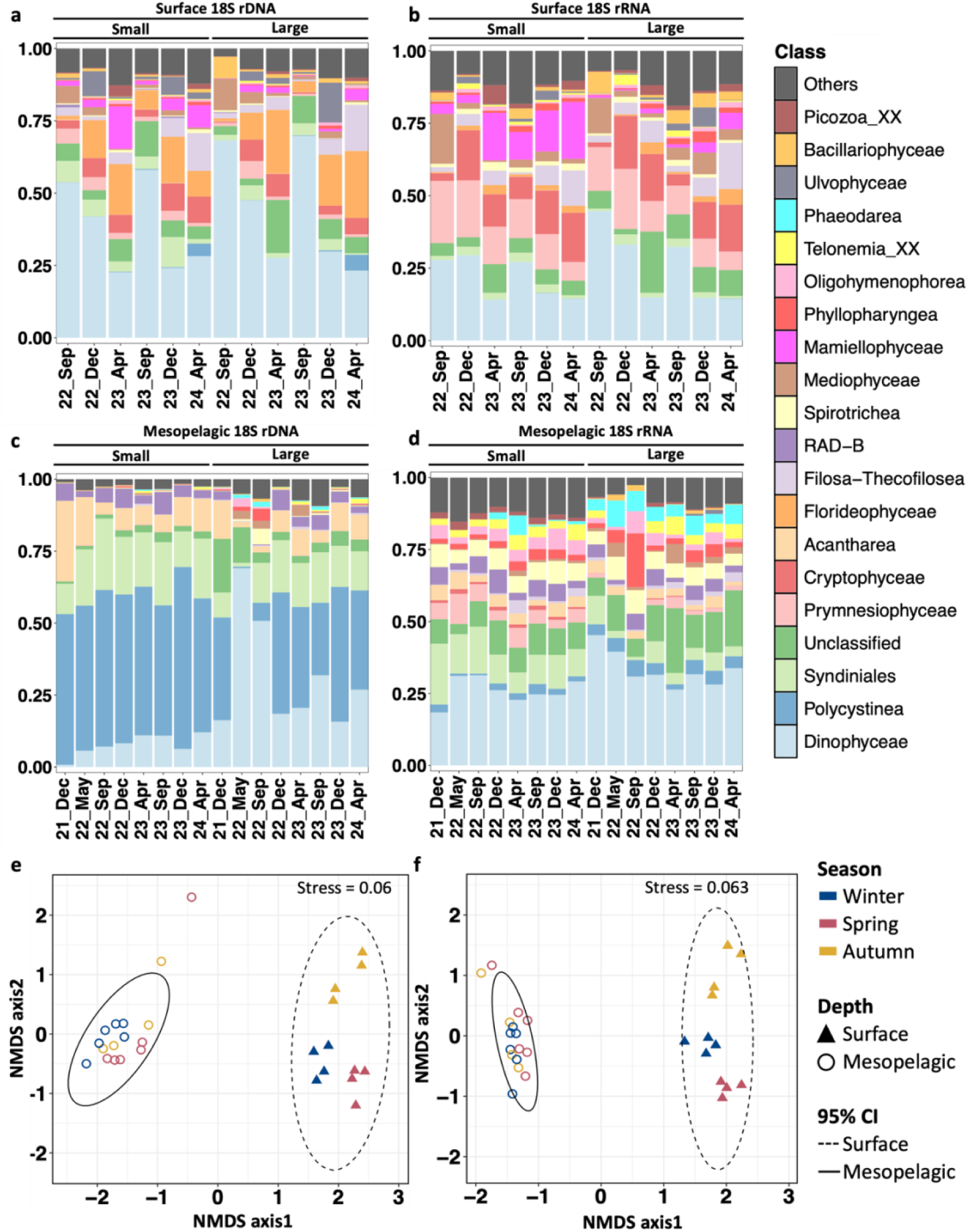
Microeukaryote community composition of the seawater samples. (a–d) Relative abundance of taxa at the class level based on the metabarcoding data for surface 18S rDNA (a), surface 18S rRNA (b), mesopelagic 18S rDNA (c), and mesopelagic 18S rRNA (d). (e,f) Non-metric multidimensional scaling (NMDS) plots showing differences in community compositions between surface and mesopelagic samples based on 18S rDNA (e) and 18S rRNA (f). 95% CI: 95% confidence interval.

To further identify microeukaryotes specially inhabiting the mesopelagic layer, we analyzed ASVs detected in the mesopelagic 18S rDNA metabarcodes and filtered out ASVs that were also detected in the surface 18S rDNA metabarcodes. The same process was also applied to the ASVs detected in the mesopelagic 18S rRNA metabarcodes. A comparison of the remaining mesopelagic-specific ASVs between the DNA and RNA metabarcodes revealed 1,094 ASVs that were detected by both 18S rDNA and rRNA metabarcodes (Fig. S2a). These ASVs were relatively abundant and apparently metabolically active, therefore representing candidate hosts for viruses inhabiting this layer. The majority (719 ASVs; 65.7%) of these 1,094 ASVs belonged to the classes Syndiniales (early branching dinoflagellates that are often parasitic), Dinophyceae, Polycystinea (silica radiolarians), Acantharea, RAD-B, and Spirotrichea (ciliates) (Fig. S2b). Dinophyceae includes a known GV host (43). We noted that one Dinophyceae ASV (ASV46) was persistently and exclusively present in mesopelagic samples, with no signal in the surface samples (Table S4). The relative abundances in different metabarcodes further revealed an important contrast between abundance and activity. Polycystinea had the highest abundance but was ranked eleventh in activity (Fig. S3d). Spirotrichea did not show high abundance but ranked third in activity (Fig. S3d). These results suggest that the mesopelagic communities are a mixture of active (living cell) and inactive (dead or dormant cell) organisms, which may shape the composition and activity of coexisting viral populations.

### GVs exclusively detected in mesopelagic metagenomes show persistency at the study site

Forty-eight GVMAGs recovered from the Muroto metagenomic data met our quality criteria (Fig. S4; see Materials and Methods), including 43 *Nucleocytoviricota* MAGs (NCV-MAGs) and 5 *Mirusviricota* MAGs (MV-MAGs) (Fig. 2a). These GVMAGs exhibited genome sizes ranging from 54 kbp to 824 kbp and GC contents ranging from 24% to 53% (Table S5). To assess the recovery level of GV lineages by GVMAGs, we investigated family-B DNA polymerase (PolB) sequences in the metagenomic dataset. Through phylogenetic analysis (Fig. S5), we identified 163 PolB sequences predicted to be of GV origin. Of these PolB sequences, 58 (36%) were found in the MAGs constructed from the metagenomic data, and 34 (21%) were found in the Muroto GVMAGs, suggesting that many GVs in the study site were not represented in our GVMAG dataset.

**Figure 2.**
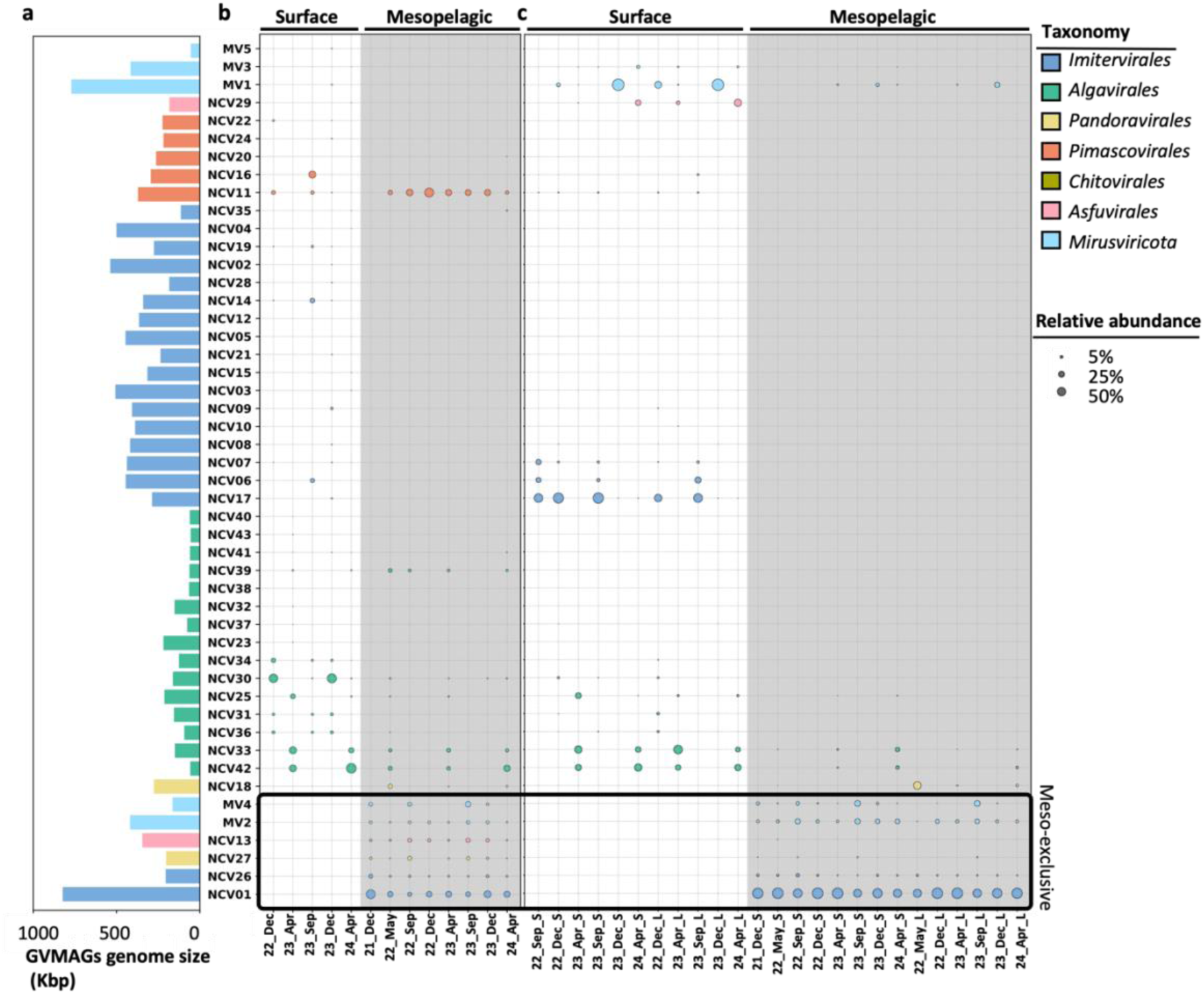
Genome size distribution and relative abundance of Muroto GVMAGs in the metagenomic and metatranscriptomic data. (a) Genome size distribution of the 48 Muroto GVMAGs. (b) Relative abundance of Muroto GVMAGs in the metagenomic data. (c) Relative abundance of Muroto GVMAGs in the metatranscriptomic data. In (b) and (c), samples from the mesopelagic layer are shown with a gray background. Circle size represents relative abundance, and circle color indicates taxonomic classification. GVMAGs enclosed by black boxes are mesopelagic-exclusive GVMAGs. X-axis labels indicate the sampling date and size fractions (S: small size fraction; L: large size fraction).

Among the 48 GVMAGs, 6 NCV-MAGs (NCV01, 13, 18, 26, 27, 42) and 2 MV-MAGs (MV2, 4) were derived from mesopelagic metagenomes, while the remaining 40 GVMAGs were assembled from surface metagenomes. Phylogenetic analysis of seven conserved NCV marker genes revealed that the six mesopelagic metagenomes derived from NCV-MAGs belonged to four orders of *Nucleocytoviricota*: two imiterviruses (NCV01, 26), one algavirus (NCV42), two pandoraviruses (NCV18, 27), and one asfuvirus (NCV13) (Fig. S6a). The two GVMAGs belonging to the order *Pandoravirales* are closely related to Emiliania huxleyi viruses (average nucleotide identify [ANI] ∼85%). The two MV-MAGs derived from mesopelagic metagenomes were classified into the MR2 marine clade (Fig. S6b) based on HK97 major capsid protein (MCP) phylogeny. Of the 40 GVMAGs derived from the surface metagenomes, 37 were NCV-MAGs, including 14 algaviruses, 17 imiterviruses, 5 pimascoviruses, and 1 asfuviruses, and 3 were MV-MAGs, including 1 in the MR2 clade and 2 in a clade composed of mirusvirus MAGs from a freshwater lake (32).

Mapping of the metagenomic reads on the 48 GVMAGs revealed that these MAGs represent 0.028%–1.28% (0.48% on average) of the total reads for the surface data compared with 0.009%–0.033% (0.020% on average) for the mesopelagic data (Fig. S7a). The lower relative abundance of GVs in the mesopelagic samples explains the lower level of GVMAG recovery from these samples. Among the 48 GVMAGs, 17 GVMAGs (15 NCV, 2 MV) were detected in the mesopelagic metagenomes, including 6 (NCV01, 13, 26, 27; MV2, 4) that were exclusively found in the mesopelagic samples (Fig. 2b, Table S6). These “meso-exclusive” GVMAGs tended to show persistent presence in the mesopelagic layer across different seasons (e.g., NCV01, NCV13, MV2) (Fig. 2b). The other 11 GVMAGs detected in the deep-sea metagenomic data were also detected in the surface metagenomes (Fig. 2b). In sharp contrast with the above six GVMAGs (i.e., meso-exclusive), many of these GVMAGs showed strong seasonal dynamics (e.g., NCV30, 42) akin to the pattern observed for GVs in a shallow coastal area (23). Endo *et al.* previously showed that the community structure of GVs in the mesopelagic layer (200–1,000 m) is different from that in the photic layer based on the *Tara* Oceans expedition data (19). Our results confirmed not only the distinct community structure but also the difference in the community dynamics between the surface and mesopelagic layers.

### Mesopelagic-specific GVs are active across seasons

Mapping of the metatranscriptomic reads on the GVMAGs revealed a similar trend to the metagenomic data (Fig. 2c, Fig. S7a,b), with the mesopelagic GV community showing seasonally stable activity and the surface layer community showing enhanced seasonal dynamics. Our observations suggest that virus–host interactions are less influenced by seasonal fluctuations in the relatively stable ecological conditions of the mesopelagic zone.

The difference in transcriptomic read mapping depth between the surface and mesopelagic layers was less pronounced than in the case of the metagenome data, with some sampling points showing relative transcriptional activity of GVMAG in the mesopelagic data comparable (or even higher) than that in the surface data. This suggests that even though fewer GVMAGs were reconstructed from the mesopelagic data, they may be highly active. For the small size fraction, the proportions of mapped transcriptomic reads were 0.0016%– 0.54% (0.11% on average) for the surface samples and 0.0024%–0.029% (0.0078% on average) for the mesopelagic samples. For the large size fraction, the proportions of mapped transcriptomic reads were 0.00028%–0.21% (0.052% on average) for the surface samples and 0.000074%–0.0017% (0.00063% on average) for the mesopelagic samples. The higher proportions of mapped transcriptomic reads in the small size fractions suggest that active infection of GVs is primarily associated with smaller host cells.

Seventeen GVMAGs showed transcriptional activity in the mesopelagic layer (Table S6). These included all six meso-exclusive GVMAGs (Fig. 2c). For five of these GVMAGs (NCV01, 26, 29; MV2, 4), transcripts were detected for over 10% of genes encoded in the genomes (Fig. S8). To the best of our knowledge, this is the first report of the transcriptional activity of GVs exclusively detected in a deep-sea environment. Furthermore, the transcriptional activities of many of these meso-exclusive GVMAGs were relatively constant over time (e.g., NCV01, NCV26, MV2, MV4) (Fig. 2c, Fig. S9). The stable community structure and activity of meso-exclusive GVs across seasons parallel the stability of the microeukaryotic community in the mesopelagic zone at the study site (Fig. 1). In contrast to the meso-exclusive GVMAGs, several GVMAGs that were detected in both surface and mesopelagic metagenomes showed transcriptional activity only in the surface samples (e.g., NCV11, NCV30). These GVs in the mesopelagic layer may have originated from the surface through sinking processes, as previously suggested (19, 29). Overall, our findings support the existence of active and stable GV populations specific to the mesopelagic zone.

### Mesopelagic GVMAGs encode specific gene functions

The six meso-exclusive GVMAGs recovered in this study contained between 51 to 913 predicted genes per genome (Table S5), with 880 genes (35.5%) being assigned to 594 KO terms (Table S7). The six meso-exclusive GVMAGs showed distinct gene compositions enriched in functions related to virus–host interactions and signaling, including components of the ubiquitin system, cytoskeletal proteins, and signal transduction regulators.

Among genes in the ubiquitin system, E3 ubiquitin–protein ligase RNF181 was found in 23 GVMAGs. Other ubiquitin system genes, such as Ariadne-1 (NCV01, NCV26, NCV27), E3 ubiquitin-protein ligase RNF13 (NCV01, NCV26), and NEDD4-binding protein 2 (NCV01, NCV26), were specific to meso-exclusive GVMAGs. Actin, myosin V, and flagellar basal-body rod protein FlgG were exclusively present in NCV01 and NCV26 and could benefit viruses by manipulating viral machinery localization during infection (14, 44, 45). Additionally, phosphoinositide 3-kinase (PI3K), a regulator of membrane dynamics and endocytic trafficking, was detected in 4 of 6 meso-exclusive GVMAGs. PI3K has been implicated in supporting the formation of replication organelles of enteroviruses and may similarly support GV replication (46).

Metatranscriptome data confirmed the active expression of these genes, with NCV01, MV2, and MV4 expressing up to 37.3%–63.6% of their genes (Fig. S8), suggesting ongoing viral replication in the mesopelagic environment. Notably, among the top 20 active KO terms in the mesopelagic samples, 7 were related to the ubiquitin system (e.g., NEDD4-binding protein 2, Ariadne-1, E3 ubiquitin-protein ligase RNF181, SIAN1, RNFT1) (Fig. S10b), compared with only 1 in the surface samples (Fig. S10a). Eukaryotic viruses can hijack ubiquitin systems helping protein folding and degradation to aid in various stages of viral propagation (47). Our results suggest that meso-exclusive GVMAGs may exploit the host ubiquitin machinery in coping with the harsh conditions (low organic carbon and light availability) in the deep ocean, as previously hypothesized for GVs in cold environments (48).

### Putative hosts of meso-exclusive GVMAGs

To gain insights into the potential hosts of the 48 GVMAGs, we performed homology searches against the Marferret v1.1.1 database. The highest homology eukaryotic matches differed between meso-exclusive GVMAGs and other GVMAGs, with the former displaying less enrichment in green algae (i.e., Mamiellophyceae and Chlorophyceae). Meso-exclusive GVMAGs were enriched in matches to Prymnesiophyceae, Dinophyceae, and Mediophyceae (Fig. S11). The taxonomic distribution of these matches was in general agreement with the microeukaryotic community structure observed by 18S metabarcoding. Regarding non-deep-sea-specific GVMAGs, MV5 and NCV18 showed unique patterns (Fig. S11). MV5 showed four genes with high homology to Cryptophyceae, which was consistent with an earlier prediction of mirusvirus hosts (6). NCV18 was found to be closely related to Emiliania huxleyi viruses, whose sequences or homologs are included in the reference *Emiliania huxleyi* (class Prymnesiophyceae) sequence data. Although these analyses imply that the hosts of meso-exclusive GVs may differ from those of other GVs, owing to limited mesopelagic protist genome representation in the current reference database, these host predictions should be regarded as preliminary. Expanding reference genomes for deep-sea microeukaryotes further is needed to improve such predictions.

### Deep-sea-specific GVs distribute widely in the global ocean

To investigate the global deep-sea-specific GV distribution, we constructed the GVGR database (see Materials and Methods) (Table S8), containing 4,473 GVMAGs and analyzed 1,890 publicly available marine metagenomes collected from depths ranging from 0 m to 10,899 m (Fig. S12). We found that GVMAG detection reached a 5,601-m depth (Fig. 3a).

**Figure 3.**
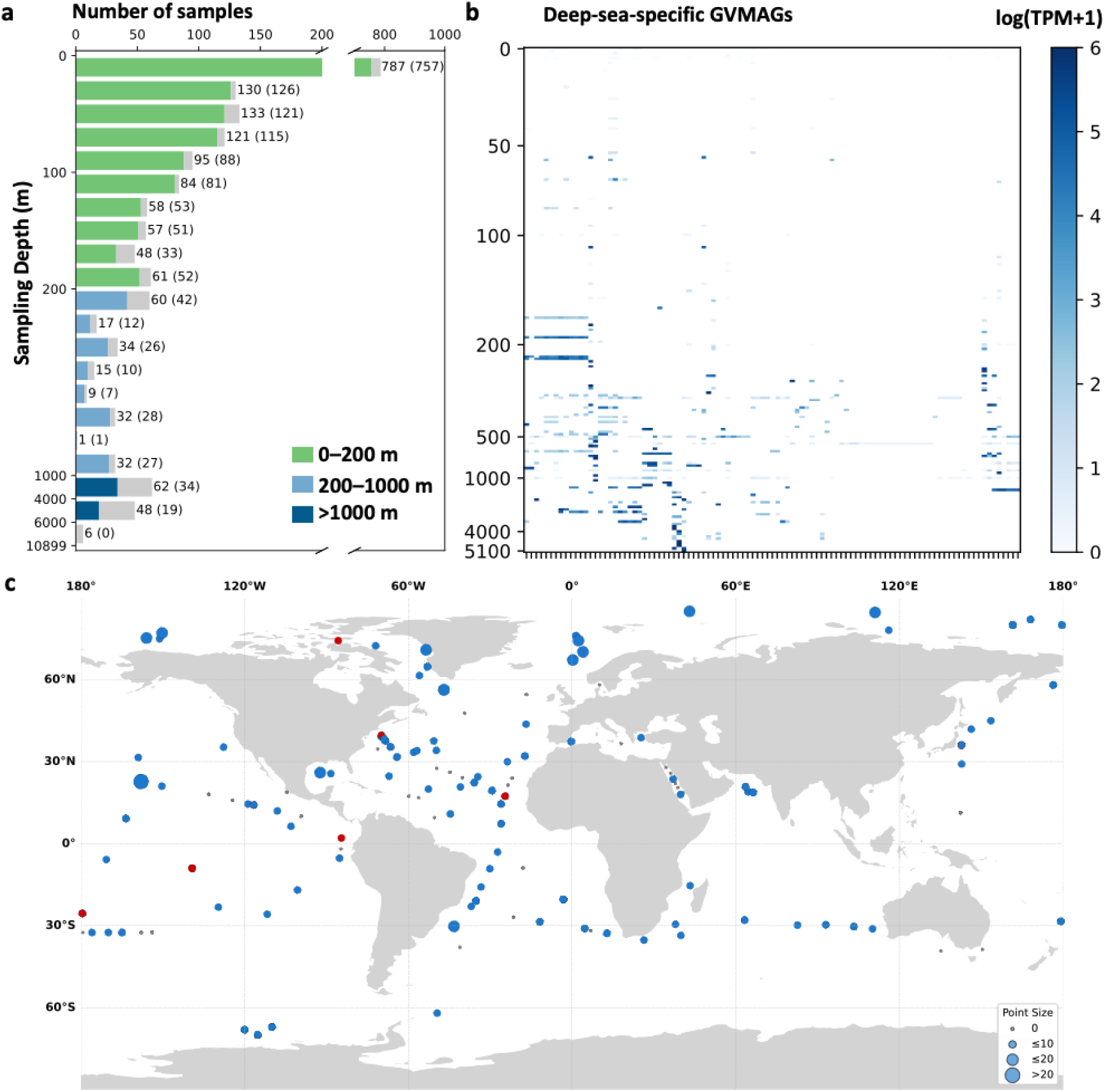
Distribution of all GVMAGs and 101 deep-sea-specific GVMAGs identified in the GVGR database across global ocean depth layers. (a) Total number of samples and number of samples with GVMAGs detected across different depths. Bars are color-coded by depth category. Numbers at the end of each bar indicate the total number of samples in that depth range; values in parentheses represent the number of samples in which GVMAGs were detected in the GVGR database. (b) Relative abundance of 101 deep-sea-specific GVMAGs identified in the GVGR database across all depth layers. Each row represents a depth, and each column represents one deep-sea-specific GVMAG, with darker blue indicating higher expression levels. (c) The deep-sea-specific GVMAG distribution across global deep-sea samples. Each point represents a single sample, with point size indicating the number of deep-sea-specific GVMAGs detected. Gray points indicate samples without deep-sea-specific GVMAGs; blue points indicate samples with deep-sea-specific GVMAGs; red points indicate samples in which ERS493705_165 (representing NCV01 after 95% sequence identity clustering) was detected.

Based on the occurrence of GVMAGs across depths, we identified 101 deep-sea-specific GVMAGs that were only or predominantly distributed in the mesopelagic or deeper layers (Fig. 3b, Table S8) (see Materials and Methods). Among them, ERS493705_165, representative of a cluster that included NCV01, exhibited a broad distribution across global deep-sea waters (Fig. 3c). Phylogenetic analysis revealed that identified deep-sea-specific GVMAGs are scattered across the tree (Fig. 4a), reminiscent of the phylogenetic distribution of GVs potentially adapted to cold environments (20). One clade within the order *Imitervirales* included seven deep-sea-specific GVMAGs and two Muroto meso-exclusive NCV-MAGs (NCV01 and NCV26). This clade was enriched with deep-sea-specific GVMAGs and showed notable high relative abundance in the deep water (Fig. 4b,c).

**Figure 4.**
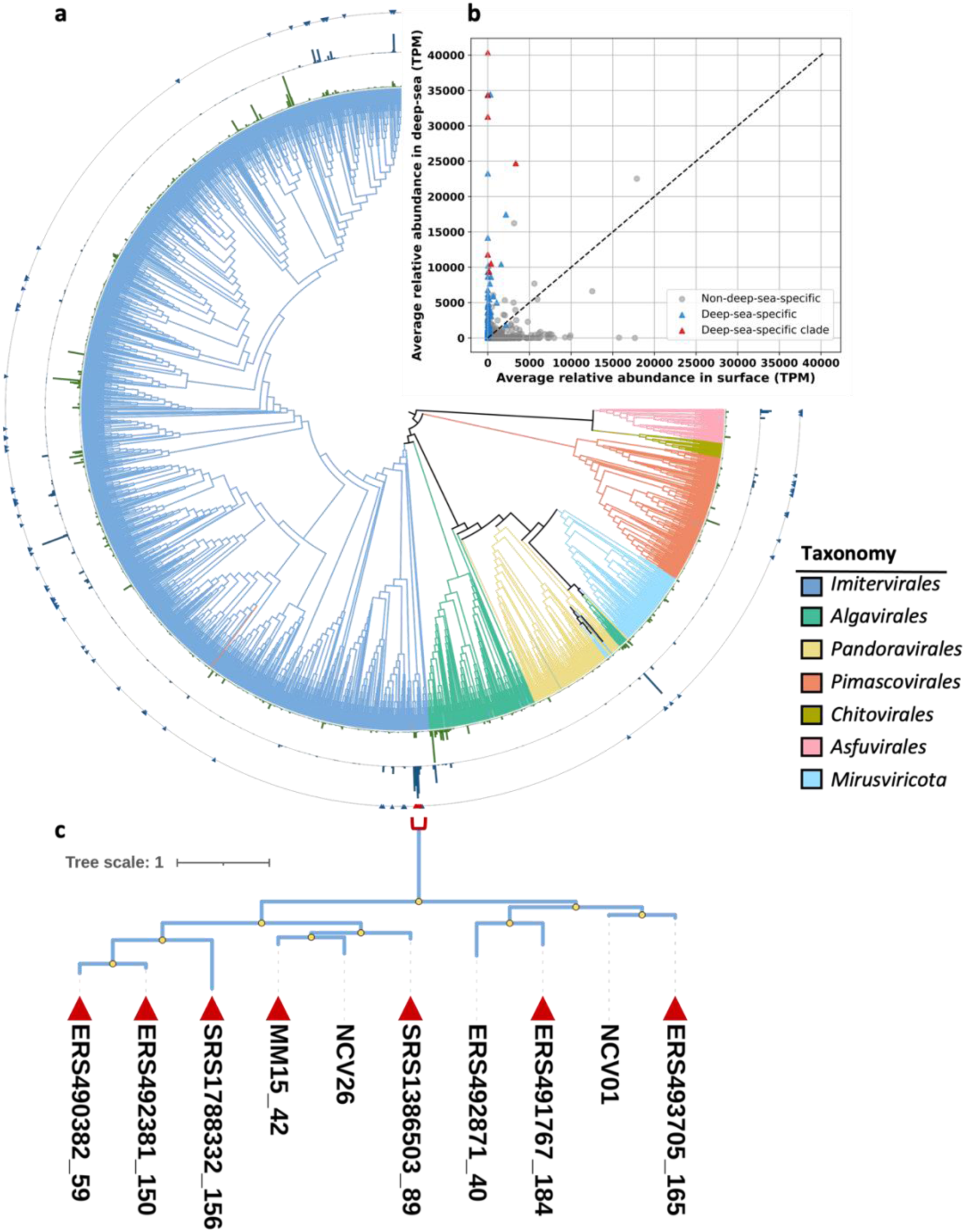
Phylogenetic diversity of GVGR genomes and deep-sea-specific GVMAGs. 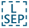 (a) Maximum-likelihood phylogenetic tree constructed from dereplicated GVGR genomes with 85% ANI, supplemented with all deep-sea-specific GVMAGs, including those derived from Muroto meso-exclusive GVMAGs. Branch colors indicate each genome’s taxonomic classification. The green bars in the inner ring indicate the average relative abundances (TPM) of the GVMAGs in photic samples. The dark blue bars in the second ring indicate the average relative abundances of the GVMAGs in deep-sea samples. Triangles on the third ring indicate deep-sea-specific GVMAGs. The red marks highlight a distinct deep-sea clade composed predominantly of deep-sea-specific GVMAGs, as described in the main text. (b) Scatterplot comparing average relative abundances of GVMAGs in surface (X-axis) and deep-sea (Y-axis) metagenomes (TPM). Each point represents one GVMAG. Red triangles indicate deep-sea-specific GVMAGs in the deep-sea clade, blue triangles denote deep-sea-specific GVMAGs, and gray circles represent other GVMAGs. The diagonal dashed line indicates equal abundance in surface and deep-sea environments. (c) Zoomed phylogenetic tree of the deep-sea clade. Branches with bootstrap support values > 95% are marked with yellow circles. The full identifier for “MM15_42” is Moniruzzaman_MM15_ERX556043_42_dc.

To investigate potential functional adaptations of deep-sea-specific GVMAGs, we identified significantly enriched KO terms and Pfam domains in deep-sea-specific GVMAGs using Fisher’s exact test with FDR correction (*P* value < 0.05) (see Materials and Methods; Fig. S13). Seventy-six KO terms were found to be enriched in deep-sea-specific GVMAGs (Table S9), including genes related to the ubiquitin system. Glutaminase and ammonium transporters were also enriched, and these functions may support energy production and nitrogen acquisition in low-energy environments (29). The above mentioned *Imitervirales* deep-sea clade showed the highest number of deep-sea-specific KO terms (Fig. S14), suggesting lineage-specific specialization. Pfam domain analysis revealed 74 domains enriched in deep-sea-specific GVMAGs (Table S9), including homologs of cytochrome P450, CTP synthases, S-adenosylmethionine decarboxylase, and a component of the iron-sulfur cluster (NifU_N). P450 has been previously reported in many GV genomes and has been proposed to be functionally connected to the 2-oxoglutarate and Fe (II)-dependent dioxygenase (49), which was also enriched in deep-sea-specific GVMAGs. The function of P450 in GVs remains unknown; however, it has been hypothesized that these genes might modulate viral or host lipid pools to aid energy production (50). Additionally, NifU is involved in Fe-S cluster formation, which play roles in diverse cellular processes. NifU has been previously reported in eukaryotes, although its specific function remains unresolved (51). Two KO terms and 22 Pfam domains were enriched in the non-deep-sea-specific GVMAGs (Table S9). These included domains for DNA damage repair (UV damage endonuclease and 5ʹ-3ʹ exonucleases), rhodopsins, phosphate transport associated proteins (PhoH), and sulfotransferases. The enrichment of PhoH suggests adaptation to phosphate-limited conditions (52, 53).

Taken together, the KO and Pfam enrichment patterns revealed that GVs in the deep-sea have evolved to encode several genes of unique function, potentially enabling them to maintain replicative efficiency by modulating host metabolic programs under environmental constraints such as limited light availability and scarce organic compounds.

## SUMMARY

Our study provided the first integrative multi-omics evidence for the persistent activity of GVs in a mesopelagic environment. Six GVMAGs from the Muroto metagenomes were exclusively detected in the mesopelagic layer, many showing persistent transcriptional activity across seasons. Some GVMAGs detected in the mesopelagic layer were active in the surface layer but inactive in the mesopelagic layer, suggesting that GV populations detected among deep-sea metagenomes may partially originate from the surface through vertical transport. Furthermore, we identified 101 deep-sea-specific GVMAGs from a global metagenomic dataset and revealed clear genomic and phylogenetic divergence between deep-sea-specific and other GVs, including identifying 76 KO terms and 74 Pfam domains enriched in the deep-sea-specific GVMAGs and phylogenetic clades enriched in deep-sea-specific GVs (Fig. 4a). Deep-sea-specific GVs exhibited distinct gene content, notably including genes for the ubiquitin system. These findings collectively support the hypothesis that distinct GV lineages may have evolved to adapt to deep-sea environments by acquiring genes to manipulate host cellular processes to thrive in the aphotic environment, which largely differs from that of the photic zone.

## MATERIALS AND METHODS

### Sample collection

Eight time-series seawater samples were collected from two depths (surface: 0.5 m, deep: 320 m) at the Kochi Prefectural Deep Seawater Laboratory, Kochi Prefecture, Japan (Fig. S1a), from December 2021 to April 2024. The seawater was pumped directly from the ocean and obtained via the deep-sea water intake system at the facility (36). The filtration system with membranes directly connected to the deep-sea water tap was built to ensure that the water remained in its original condition (Fig. S1b, Table S2). After pre-filtration through a 150-µm-pore-size nylon mesh to remove large organisms, a large volume of seawater, especially for deep-sea samples (30–1110 L), was filtered through 3-µm- or 5-µm-pore-size polycarbonate membranes (Merck, Germany) and 0.2-µm-pore-size Sterivex-GP PES filters (Merck). This filtration step was designed to collect large (3–150 µm or 5–150 µm) and small (0.2–3 µm, 0.2–5 µm, or 0.2–150 µm) size fractions. All filtrated membranes were immediately frozen and transported in a cryo-shipper at liquid nitrogen temperatures to the laboratory and stored at −80°C until DNA and RNA were extracted. Filtration and cryo-preservation of each filter were completed within 25 minutes.

Total DNA and RNA were extracted using the AllPrep RNA/DNA kit (Qiagen, Germany) following the protocol described by Okazaki *et al.* (54). The DNA/RNA quantity was assessed using a Qubit fluorometer (Thermo Fisher Scientific, USA). The extracted DNA/RNA was used for 18S rDNA metabarcoding (28 samples), 18S rRNA metabarcoding (28 samples), metagenomic (13 samples), and metatranscriptomic (25 samples) sequencing (Fig. S1c). Environmental variables, including temperature, salinity, and pH were recorded continuously by the Kochi Prefectural Deep Seawater Laboratory (Table S10).

### Metabarcoding sequencing and analyses

Microeukaryote amplicon sequencing was performed for both DNA and RNA-derived cDNA (rDNA and rRNA sets, respectively). For RNA, genomic DNA removal and first-strand cDNA synthesis were conducted using the SuperScript IV VILO Master Mix with the ezDNase enzyme kit (Thermo Fisher Scientific) following the manufacturer’s protocol. The V4 region of the 18S rRNA gene from both cDNA and DNA was amplified by the KAPA HiFi HotStart ReadyMix (Roche) using the universal eukaryotic primer set E572F/E1009R (55) attaching Illumina overhang adapters. The PCR conditions were as follows: initial pre-denaturation at 95°C for 2 minutes, followed by 30 cycles of denaturation at 98°C for 20 seconds, annealing at 61°C for 15 seconds, and extension at 72°C for 30 seconds, with a final extension at 72°C for 2 minutes. Triplicate PCR products were pooled and then purified using VAHTS DNA Clean Beads (Vazyme). The amplicon libraries were sequenced on the Illumina MiSeq platform to generate paired-end reads (2 × 300 bp).

Raw sequencing reads were processed using QIIME2 v2024.10 (56) with the DADA2 plugin (57). The “dada2 denoise-paired” command was used for adapter removal, primer removal, low-quality read trimming, dereplication, chimera removal, and amplicon sequencing variant (ASV) identification. Rare ASVs that appeared in less than two sequences across all samples were excluded. Taxonomic classification was performed on the remaining ASVs based on a pre-trained naive Bayes classifier trained against the PR2 reference database v5.0.0 using the “feature-classifier classify-sklearn” plugin (58). ASVs classified as non-protist lineages (e.g., metazoa, fungi) were removed. Finally, 1,616,977 reads from 56 samples were grouped into 5,031 ASVs. To enable comparisons between samples, the ASV table was rarefied to the minimum read depth per sample (7,581 reads), resulting in a final dataset of 4,981 ASVs and 424,536 reads. Statistical differences among sample groups (e.g., depth and season) were analyzed using PERMANOVA with 9,999 permutations based on the Bray–Curtis dissimilarity calculated from the Hellinger-transformed ASV abundance table. All statistical analyses were conducted using the Vegan v2.6.10 package in the R software (59).

### Metagenomic sequencing and analyses

Metagenomic shotgun sequencing libraries were constructed using the Illumina DNA Prep kit (Illumina) and sequenced on the Illumina NovaSeq 6000 platform, generating 150-bp paired-end reads with an average of 58 Gbp per sample. Raw reads were quality-controlled using fastp v0.23.4 (60) with default parameters. For each metagenome, trimmed reads were assembled into contigs using MEGAHIT v1.2.9 (61) in “meta-sensitive” mode. Contigs over 2.5 kb were retained for downstream analysis. Trimmed reads from all samples were primarily used for cross-mapping against contigs assembled from individual samples by Bowtie2 v2.4.5 (62). Additionally, a co-assembly was performed by pooling reads from all deep-sea samples. Contigs were clustered into bins using MetaBAT2 v2.15.15 (63). Metagenomic bins were generated for each of the 13 metagenomes and for the pooled deep-sea metagenomes. Genes in MAGs were predicted using Prodigal-gv v2.11.0 (64, 65) with “meta” mode for marker gene identification and functional annotation.

NCV-MAGs and MV-MAGs were reconstructed separately following previously published methods (23, 32). To identify NCV-MAGs, we used a custom pipeline named hedera v.0.0.5 (https://github.com/banhbio/hedera). Briefly, the pipeline employs a gene density index [NCLDV index (23)] calculated using the genome size and the presence of 20 marker genes selected from the Nucleo-Cytoplasmic Virus Orthologous Groups (NCVOG) (66). MAGs with an NCLDV index over 5.75 are identified as potential NCV-MAGs. The NCV verification and decontamination is performed using the results of Viralrecall v2.1 (score > 0) (67), Virsorter2 v2.2.3 (max score group “NCLDV”) (68), CAT v5.2.3 (“Nucleocytoviricota”) (69), and hidden Markov models (HMMs) built with 149 NCVOGs (E-value < 1.0×10^−3^) (23). Bins selected by hedera were further investigated for the possibility of chimeric bins (e.g., multiple viral genomes) by manual inspection using Anvi’o v8 (70). First, we primarily focused on deep branching clades that showed markedly different occurrence patterns or GC contents from other clades. If such clades showed the seven core marker genes of GVs (PolB, A32 packaging enzyme, superfamily II helicase, VLTF3 transcriptional factor, topoisomerase family II, transcription factor IIB, and RNA polymerase second largest subunit [RNAPS]), they were retained as a different bin, otherwise, they were discarded. Second, for the remaining clades, if we identified clearly different patterns of occurrence or GC content, we split the bins into two or more sub-bins. Finally, the modified bins were re-assessed for the NCLDV index and only those with an NCLDV index over 5.75 were retained. For MV-MAGs, bins were identified as a mirusvirus if the HK97 MCP gene was detected using HMMER v3.4 (E-value < 1.0×10^−3^, bit score > 100) (71). After manual inspection with Anvi’o, MAGs containing a contig encoding HK97-MCP were retained. Decontamination was performed using CheckV v1.0.1 (database v1.4) (72) by concatenating all contigs of each MAG and using “end_to_end” mode to remove prokaryotic contigs and provirus segments (32). Dereplication was conducted using dRep v3.5.0 (73) at an average nucleotide identify (ANI) of 95% with the parameters “--ignoreGenomeQuality -- S_algorithm ANImf -sa 0.95 --clusterAg single -sizeW 1”. The resulting non-redundant Muroto GVMAGs (NCV-MAGs and MV-MAGs) were species-level representatives. GVMAG names were assigned as follows: “NCV” or “MV”, followed by a serial number indicating the rank of genome size from large to small (e.g., NCV10, MV5).

To determine the relative abundances of Muroto GVMAGs, the coverage and transcripts per million (TPM) (74) of each MAG were calculated by mapping all quality-controlled trimmed reads to Muroto GVMAGs using CoverM v0.6.1 (75).

### Metatranscriptomic sequencing and analyses

RNA samples of both the large and small size fractions from the surface and deep-sea were subjected to metatranscriptomic sequencing. Poly-A strand-specific RNA sequencing libraries were constructed using the NEBNext® Poly(A) mRNA Magnetic Isolation Module and NEBNext® Ultra ll Directional RNA Library Prep Kit (New England Biolabs, USA). Libraries were sequenced on the Illumina NovaSeq 6000 platform and 150-bp paired-end reads were generated. An average of 24 Gbp of sequence per sample was obtained. Raw reads were quality-controlled using fastp v0.23.4 with the “-l 100” parameter to remove low-quality reads and reads shorter than 100 bp. SortMeRNA v4.3.6 (76) was used to filter out rRNA reads against the Silva v138.1 LSU NR99, Silva v138.1 SSU NR99, RFAM 5s, and RFAM 5.8s databases using default settings. Filtered mRNA reads were mapped against the Muroto GVMAGs using Salmon v1.10.1 (77) with the parameters “--meta -- minScoreFraction 0.95 --validateMappings”. Transcript abundance was normalized as TPM. The results were used to quantify the expression abundance of each gene and MAG.

### Compiling the GVGR database and identifying deep-sea-specific GVMAGs

To investigate the global deep-sea GV distribution and adaptation, we compiled a comprehensive giant virus genome reference, named GVGR, by integrating the Muroto GVMAGs with publicly available GV metagenomic datasets (Fig. S4). We first extracted NCV-MAGs from metagenomic bins generated using MetaBAT2 by the OceanDNA MAG project (38). Metagenomic bins with an NCLDV index over 5.75 were considered as potential NCV-MAGs. These MAGs were decontaminated through retaining contigs identified as “NCLDV” by Virsorter2 v2.2.4 or those with a ViralRecall v2.1 score greater than 0. Host-derived sequences and proviral segments were removed using CheckV v1.0.1, as mentioned above. For MV-MAG classification, the initial screening was consistent with the process described for Muroto MV-MAGs. Contigs classified as “chromosome” or “plasmid” using geNomad v1.8.0 (65) were excluded. MAGs with a size between 50 kbp–3 Mbp in length were retained as GVMAGs.

Additionally, we incorporated publicly available GV genomes from multiple sources, including GOEVdb (2), 1,065 GVMAGs from Uranouchi Inlet, Japan (23), 293 LBGVMAGs from Lake Biwa (32), and four circular endogenous *Mirusviricota* (6, 39). All collected MAGs were pooled and dereplicated using dRep v3.5 at an ANI of 95% as previously mentioned. Within the ANI-based cluster, the MAGs with the highest N50 value among the top 30% largest sized genomes were selected as representative.

To assess MAG quality, a phylogeny-informed MAG assessment (PIMA) based on a phylogenetic tree and orthologous groups (OGs) was performed on the above-mentioned dataset and Muroto GVMAGs (23). For phylogenetic tree reconstruction, seven conserved marker genes were identified using the script “ncldv_markersearch.py”(78). For MV-MAGs, four marker genes (HK97-MCP, RNAPS, RNA polymerase largest subunit [RNAPL], and PolB) were identified using HMMER v3.4 (71) with pairwise alignment against the reference HMM profiles. These marker genes were aligned and concatenated using Clustal Omega v1.2.4 (79), trimmed with trimAl v1.2.1 (-gt 0.1) (80) and a phylogenetic tree was built with FastTree v2.1.11 (81) using default parameters, serving as an input for PIMA of NCV-MAGs and MV-MAGs, separately. OGs were classified using GVOG HMM profiles (82) using HMMER v3.4 (E-value < 1.0×10^−10^) (71). If multiple OGs were assigned to a single sequence, the OG with the lowest E-value was retained.

PIMA was applied at a relative evolutionary divergence threshold of 0.65, corresponding to genus- or family-level classification (23). MAG consistency was evaluated based on the proportion of core genes present, while redundancy was calculated as the proportion of duplicated core genes exceeding the mode copy number observed across all MAGs within each lineage. MAGs exhibiting redundancy over 50% were removed. A second round of dRep dereplication was subsequently performed to integrate Muroto GVMAGs, producing the final GVGR dataset. The taxonomy of GVMAGs in the GVGR dataset was validated using TIGTOG (83). Gene prediction and abundance analyses in all 1,890 global samples (38) were performed using methods identical to those described above.

Each of these GVMAGs was categorized as “deep-sea-specific” or “non-deep-sea-specific”. We considered genomes showing overrepresentation in deep-sea samples (depth > 200 m) as deep-sea-specific GVMAGs. The overrepresentation was ascertained using Mann– Whitney U tests with Benjamini–Hochberg (BH) corrected *P* values (*P* < 0.05) (84). We also considered genomes with signals only in deep-sea samples (TPM > 0) as deep-sea-specific GVMAGs. Other GVMAGs, not assigned to deep-sea-specific category, were referred to as “non-deep-sea-specific”.

### Protein prediction and annotation

Proteins predicted from Muroto GVMAGs and the GVGR database were annotated with KEGG orthology (KO) using KofamScan v1.3.0 (85). KO annotations with the lowest E-values were retained. We further generated Pfam annotations of the genomes using AnnoMazing (https://github.com/BenMinch/AnnoMazing), a pipeline to annotate proteins based on HMM profiles. The annotations were performed using the Pfam database (86) with an E-value cut-off of 1.0×10^−5^.

To identify KO terms and Pfam domains significantly enriched in deep-sea-specific GVMAGs, Fisher’s exact test was conducted to compare the frequency of each annotated KO term in deep-sea-specific GVMAGs and non-deep-sea specific GVMAGs. KO terms with a total frequency ≤ 2 were excluded from further analysis.

To infer the potential eukaryotic hosts of GVs, we performed protein homology searches using BlastP in Diamond v2.1.10 (E-value cut-off of 1.0×10^−5^ and bit score > 50) (87) against the MarFERReT ver1.1.1 database (88). Hits with the lowest E-values were retained and used to assign candidate host taxonomy.

### Analyses of phylogenetic diversity and clade delineation

A phylogenetic tree was first constructed based on PolB sequences (>500 amino acids) extracted from Muroto contigs (>1 kbp). The GOEV database and a wide range of eukaryotic and additional viral lineages (2, 89) were used as a reference set of PolB sequences to evaluate the completeness of GV recovery in Muroto GVMAGs. Another tree for *Mirusviricota* was generated using HK97-MCP sequences from Muroto MV-MAGs, combined with previously reported *Mirusviricota* genomes from the GOEV database, LBGVMAGs, and circular endogenous MV-MAGs. For *Nucleocytoviricota*, a concatenated phylogenetic tree was constructed using the seven core marker genes (82) for Muroto NCV-MAGs, along with 220 reference genomes from culture (2). Another concatenated phylogenetic tree was generated using the same seven core marker genes, incorporating all deep-sea-specific MAGs, Muroto meso-exclusive MAGs, and the GVGR dataset redundant by dRep v3.5.0 at an ANI of 85%. The marker gene search, alignment, concatenation, and trimming methods were the same as those described for the PIMA above. The tree was inferred using IQ-TREE v2.2.0 (90) with the ModelFinder Plus option to determine the best-fitting model. Support values were inferred using 1,000 ultrafast bootstraps (91). Tree visualization and rooting were carried out using iTOL v7 (92).

## ACKNOWLEDGEMENTS

We thank the Kochi Prefectural Deep Seawater Laboratory for providing sampling permission and environmental metadata. We thank Hiroto Sasaki, Jun Xia, Qingwei Yang, Pascal Hingamap, and Keizo Nagasaki for their valuable assistance with sampling. Computational work was supported by the SuperComputer System, Institute for Chemical Research, Kyoto University. We thank Edanz (https://jp.edanz.com/ac) for editing a draft of this manuscript.

W.L. led the sampling, performed all experiments and most of the bioinformatics analyses. Y.N., J.W., K.N., and W.L. contributed to the compilation of the GVGR database. K.N. together with W.L. performed the statistical analysis on the GVMAGs from the GVGR database. H.B. contributed to the construction of the NCV-MAG pipeline. E.M. and L.M. contributed to part of the bioinformatics analyses. M.M. supervised E.M. T.Y. initiated the use of the Kochi Prefectural Deep Seawater Laboratory for this sampling. R.Y.N. contributed to the design of filtration systems. H.O. conceived the study and supervised W.L. and K.N. H.E. and Y.O. co-supervised W.L. and K.N., and participated in sampling. W.L. generated the initial draft of the manuscript. All authors contributed to the interpretation of data and writing of the manuscript, and all approved the final draft.

## DATA AVAILABILITY

The data generated from the article, the processed data, and metadata are available in the DNA database of Japan (DDBI) at www.ddbj.nig.ac.jp, and can be accessed under accession numbers PRJDB35423 and PRJDB35424. The nucleotide and protein sequences of the Muroto GVMAGs and GVGR database constructed in this study are available at GenomeNet: https://www.genome.jp/ftp/db/community/GVGR_MurotoGV. Other data used in this study include the Giant Virus Orthologous Groups (GVOGs) database (https://faylward.github.io/GVDB/).

## SUPPLEMENTAL MATERIAL

### Supplemental Tables

Table S1: Summary of sequence data obtained from the Muroto samples

Table S2: Sampling methods used in this study

Table S3: Total extracted DNA and RNA from the Muroto samples collected from the surface and mesopelagic layers

Table S4: ASV abundance table based on the Muroto metabarcoding sequencing data

Table S5: Information on the 48 Muroto GVMAGs

Table S6: Relative abundance of the 48 Muroto GVMAGs in the metagenomic and metatranscriptomic samples

Table S7: KO annotations for the 48 Muroto GVMAGs

Table S8: Information on the GVGR database

Table S9: Deep-sea-specific and non-deep-sea-specific KO terms and Pfam domains identified from the GVGR database

Table S10: Environmental parameters measured at the Kochi Prefectural Deep Seawater Laboratory

## Supplemental Figures

**Figure S1.**
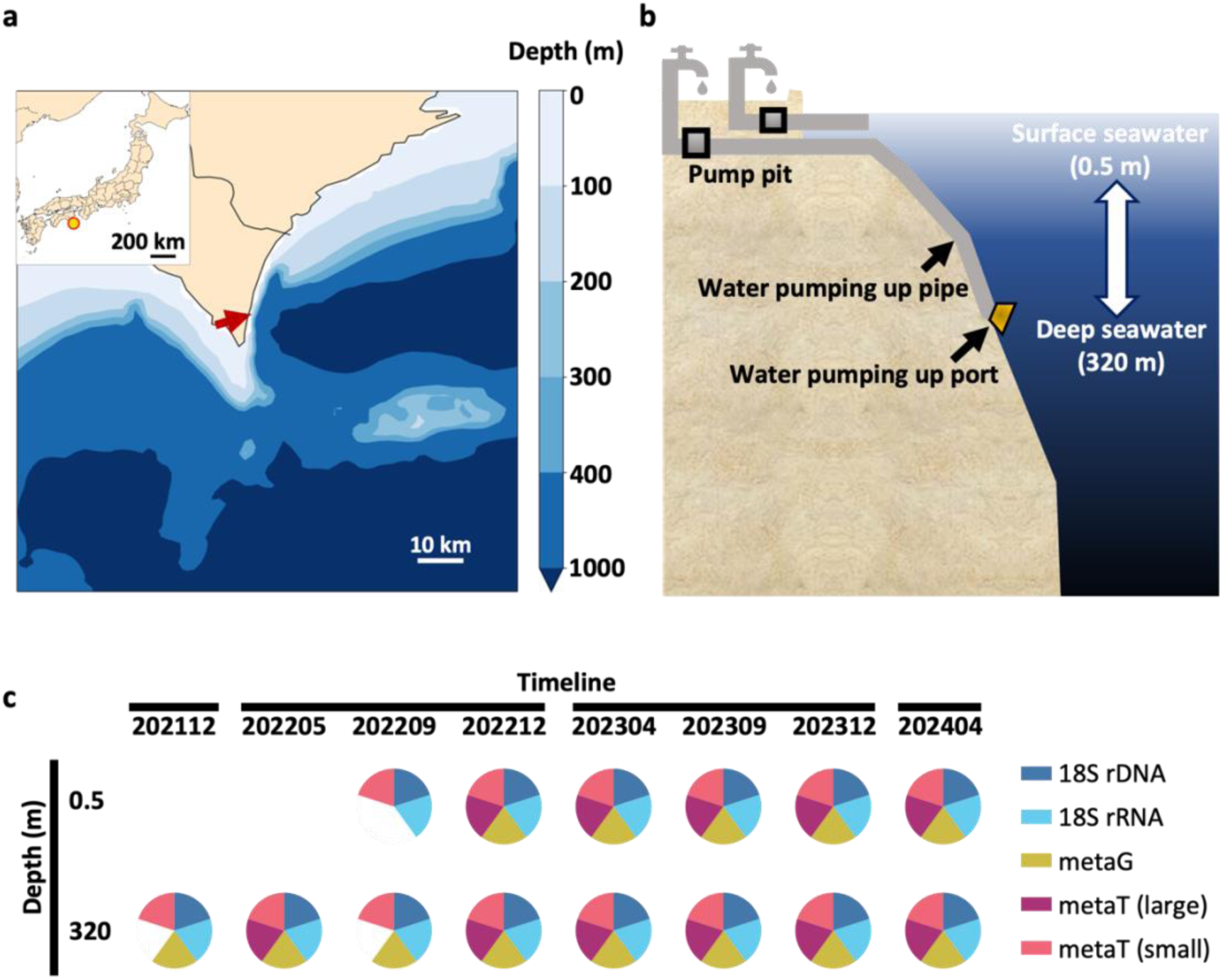
Sampling location and depth. (a) Location of the sampling site at the Kochi Prefectural Deep Seawater Laboratory. (b) Schematic illustration of surface and deep seawater intake equipment. (c) Diagram showing the sampling year and month as well as depth, either surface (0.5 m) or deep-sea (320 m). 18S rDNA: DNA-based 18S rRNA metabarcoding, 18S rRNA: cDNA-based 18S rRNA metabarcoding, metaG: metagenomic sequencing, metaT (small): metatranscriptomic sequencing for small size fraction, metaT (large): metatranscriptomic sequencing for large size fraction.

**Figure S2.**
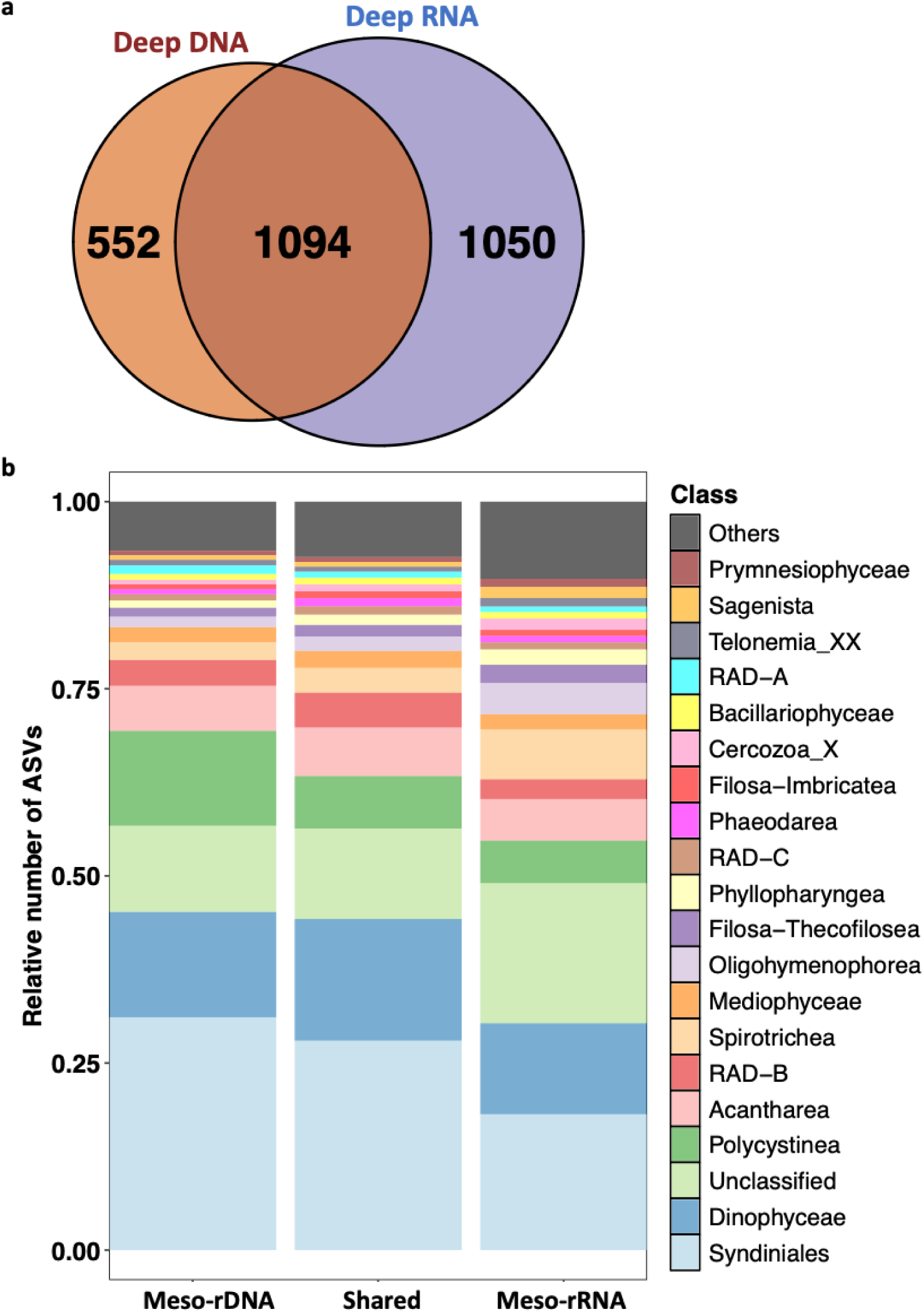
Composition of deep-sea-specific ASVs. (a) Venn diagram showing the overlap of taxa between deep-sea 18S rDNA and 18S rRNA after discarding ASVs detected in surface 18S rDNA and 18S rRNA, respectively. (b) Class-level taxonomic composition of microeukaryotes based on the number of ASVs in three categories: Deep DNA, Shared ASVs, and Deep RNA. Only the top 20 classes (based on the number of ASVs within the Shared group) are shown in the plot. Taxonomic classes outside the top 20 were grouped into “Others” to simplify visualization. “Unclassified” refers to ASVs that could not be taxonomically assigned at the class level owing to insufficient reference database annotation.

**Figure S3.**
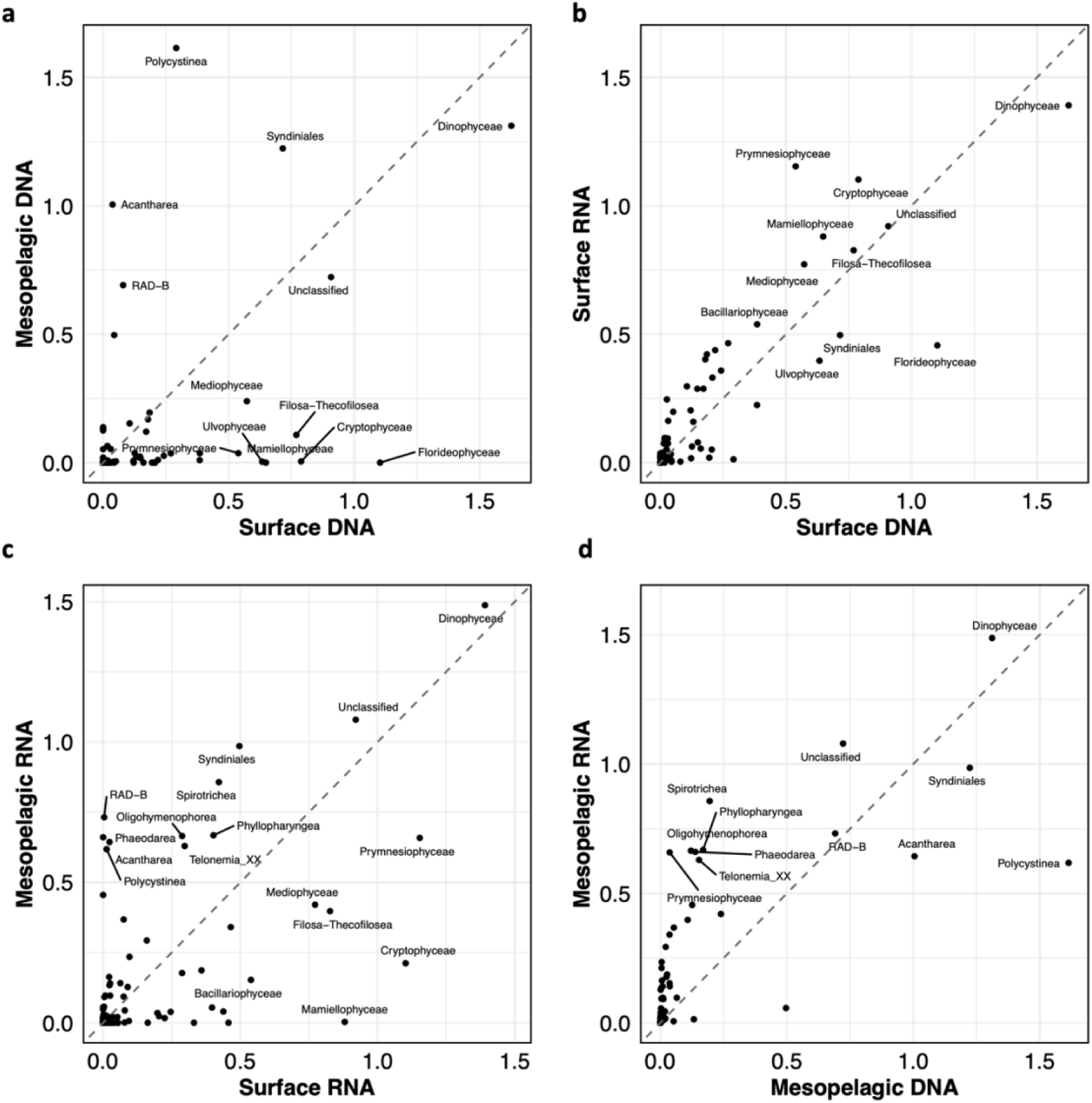
Comparisons of mean relative abundance (logTPM+1) of taxa at class level classification based on 18S rDNA/rRNA metabarcoding data. The taxa with logTPM+1 > 0.5 are labeled.

**Figure S4.**
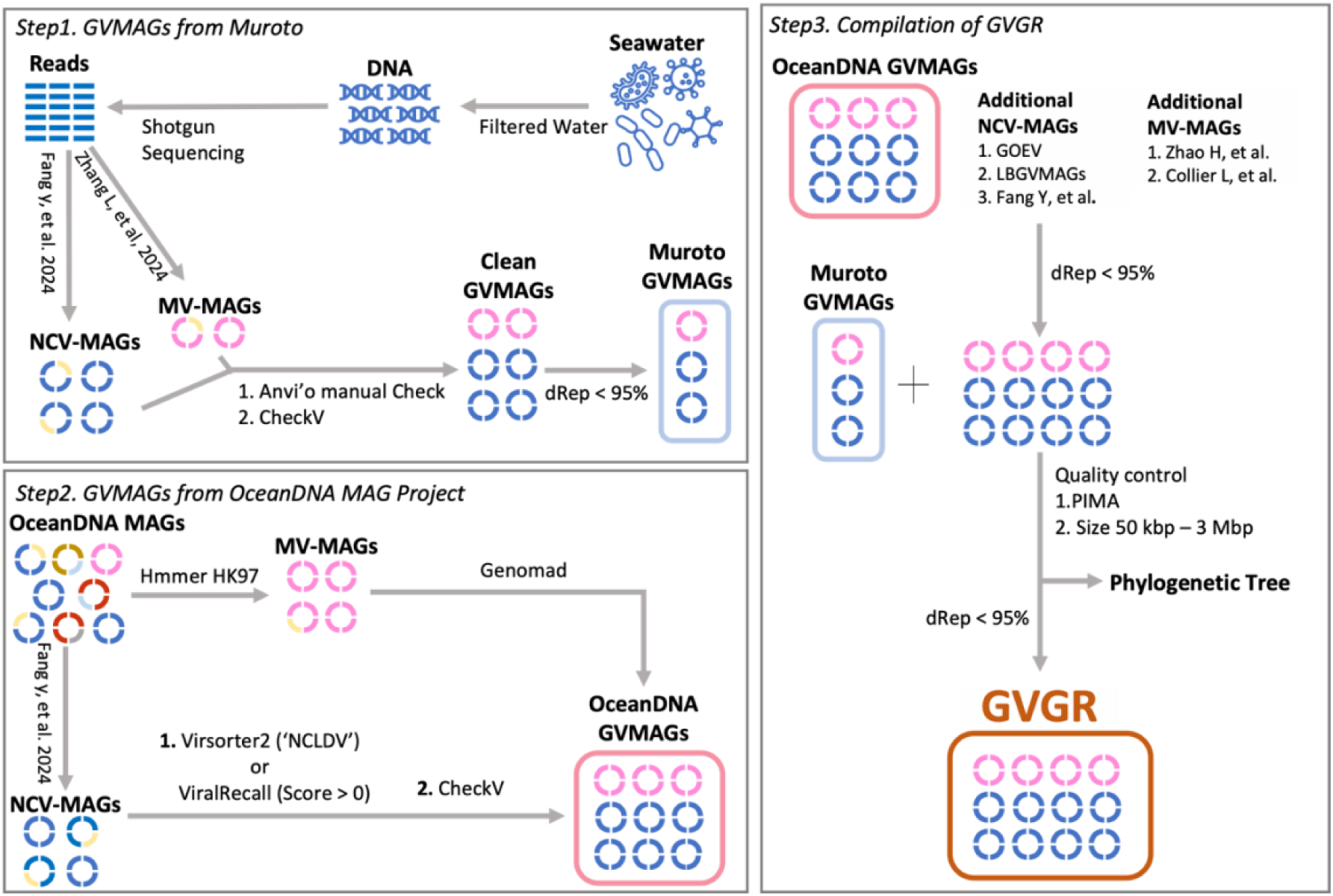
Giant virus genome reference (GVGR) database. The pipeline for the construction of the GVGR database is composed of three steps. Step 1 generates manually curated GVMAGs (Muroto GVMAGs) from metagenomic data generated in this study. Step 2 extracts GVMAGs (named OceanDNA GVMAGs) from the OceanDNA MAG project. Step 3 combines Muroto GVMAGs, OceanDNA GVMAGs, and previously published GVMAG datasets to generate a comprehensive giant virus genome reference (GVGR) database.

**Figure S5.**
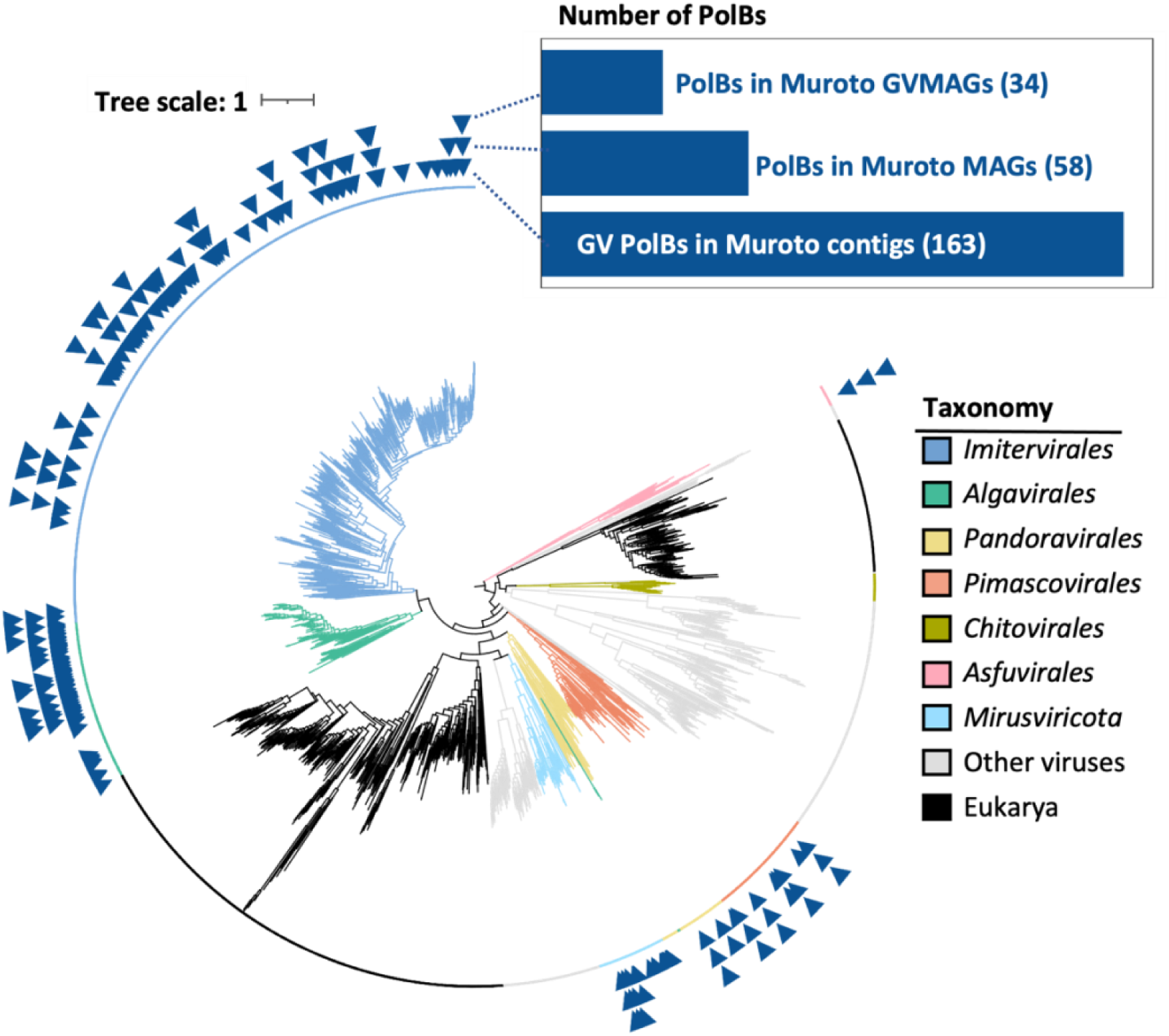
Phylogenetic analysis of PolB from the Muroto samples. Maximum-likelihood phylogenetic tree of PolB from Muroto metagenomic contigs (>2.5 kbp) and reference sequences (Gaïa et al., 2023; Kazlauskas et al., 2020). The colors of the branches and the most inner ring indicate viral classification. The three outer rings with triangles indicate the distribution of PolB sequences: the first ring shows PolB sequences likely to be of GV origin and found in the Muroto contigs (>2.5 kbp), the second ring shows PolB sequences likely to be of GV origin and found in Muroto MAGs, and the outermost ring indicates PolB sequences in the qualified Muroto GVMAGs. The bar plot indicates the number of PloB sequences in each ring.

**Figure S6.**
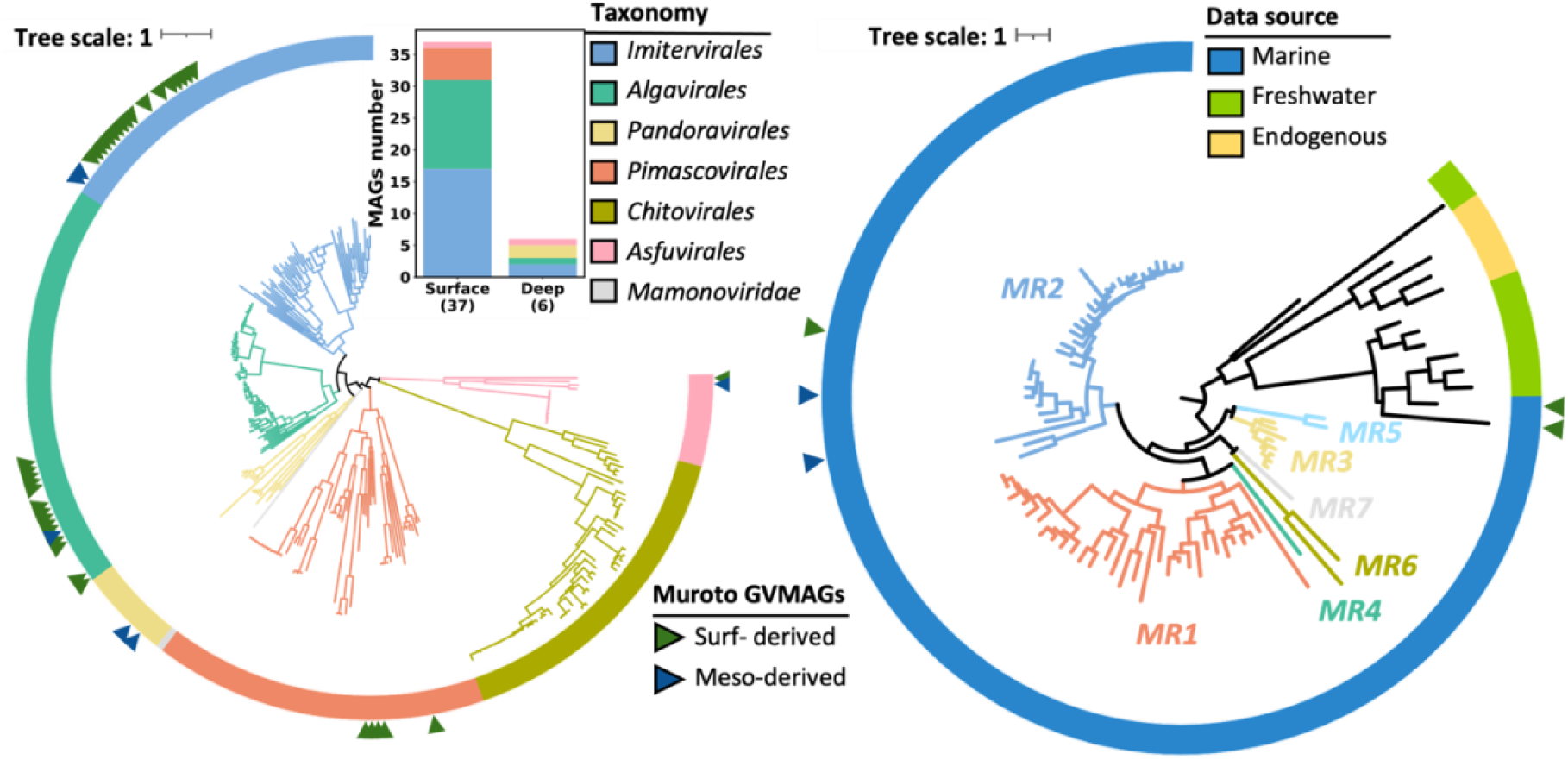
Phylogenetic and taxonomic composition of Muroto GVMAGs. (a) Maximum-likelihood phylogenetic tree of Muroto NCV-MAGs and 220 reference genomes inferred using IQ-TREE, based on seven conserved marker genes. Ring and branch colors indicate taxonomic classifications. (b) Phylogenetic tree of Muroto MV-MAGs and reference genomes inferred using IQ-TREE, based on the HK97 MCP gene. The ring indicates the MAG source, while branch colors represent clade classifications. “MR” marks the mirusvirus-associated clade, and its label color corresponds to the color of the associated branch. Blue and green triangles denote deep-sea and surface-derived GVMAGs, respectively.

**Figure S7.**
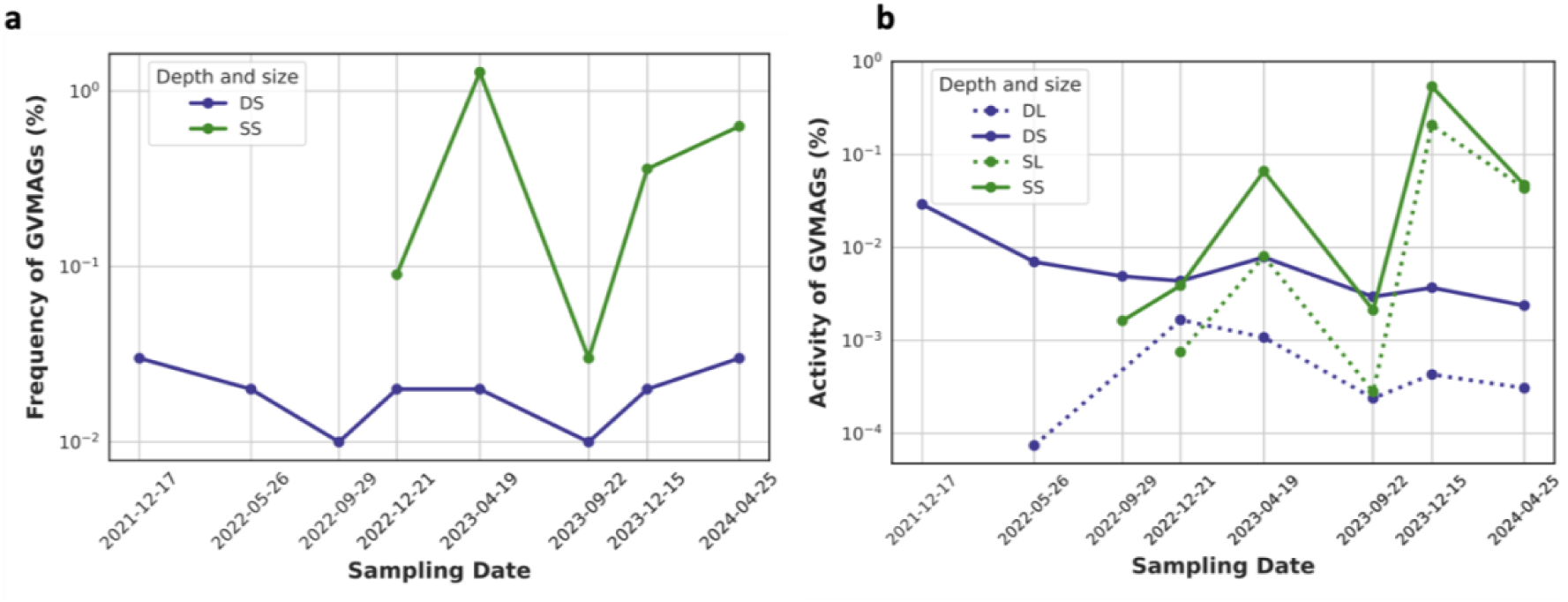
Dynamics of the presence and activity of Muroto GVMAGs. (a) Percentage of metagenomic reads mapped to the 48 Muroto GVMAGs over time. (b) Percentage of metatranscriptomic reads mapped to the 48 Muroto GVMAGs over time. Blue lines represent deep-sea samples, while green lines represent surface samples. Solid lines indicate small-size fractions (0.2–3 µm, 0.2–5 µm, or 0.2–150 µm), whereas dashed lines correspond to large-size fractions (3–150 µm or 5–150 µm). Time points are plotted based on actual sampling intervals, and the spacing on the X-axis reflects real temporal distances.

**Figure S8.**
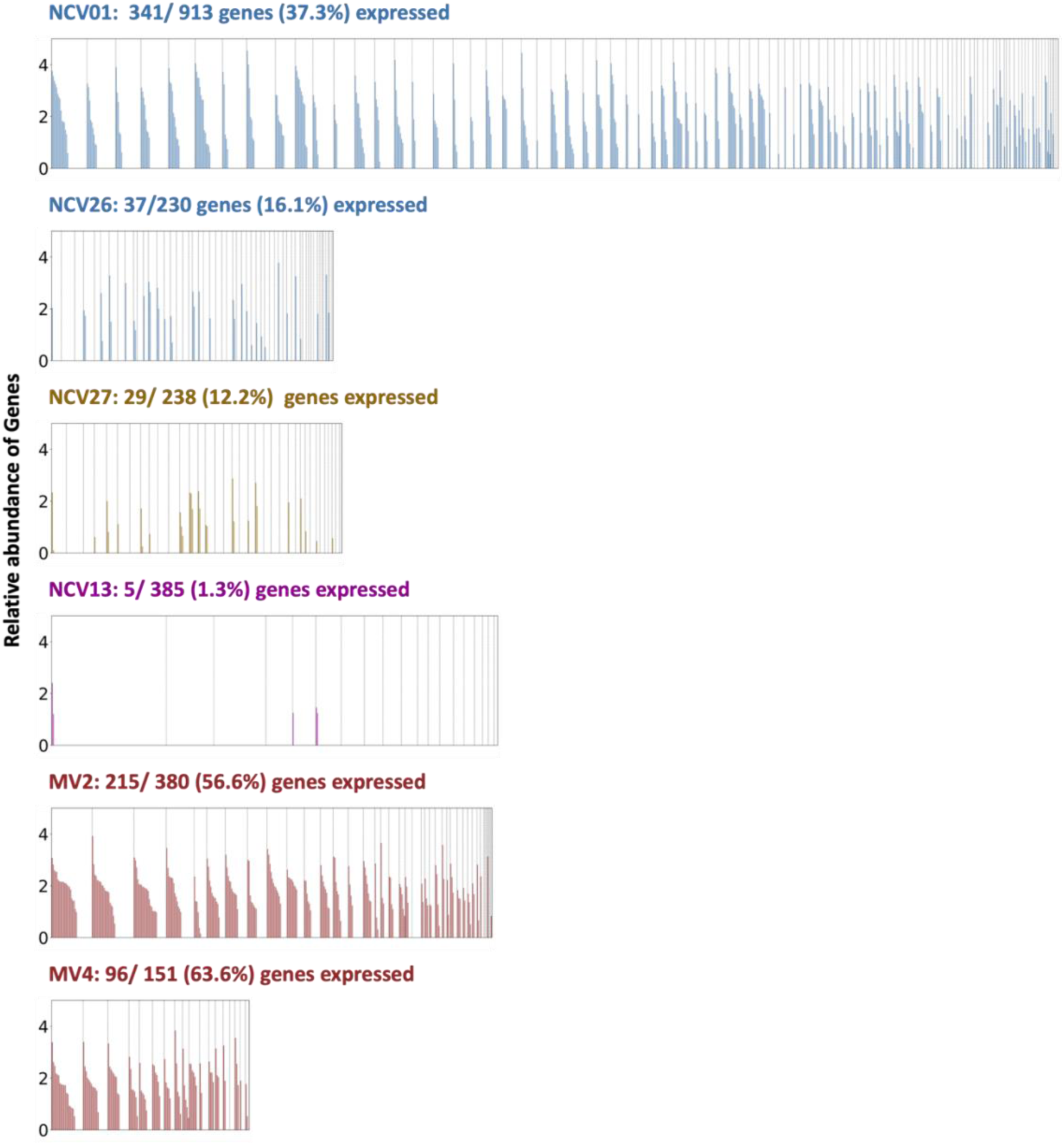
Gene expression profiles of six meso-exclusive GVMAGs. Each horizontal panel represents a meso-exclusive GVMAG, with the X-axis showing individual genes and the Y-axis indicating mean relative levels (logTPM+1). Bar colors correspond to GV classifications. Vertical dashed lines indicate the boundary between contigs. Within individual contigs, genes are ordered by their expression levels. The total number of genes and the percentage of expressed genes are annotated for each GVMAG.

**Figure S9.**
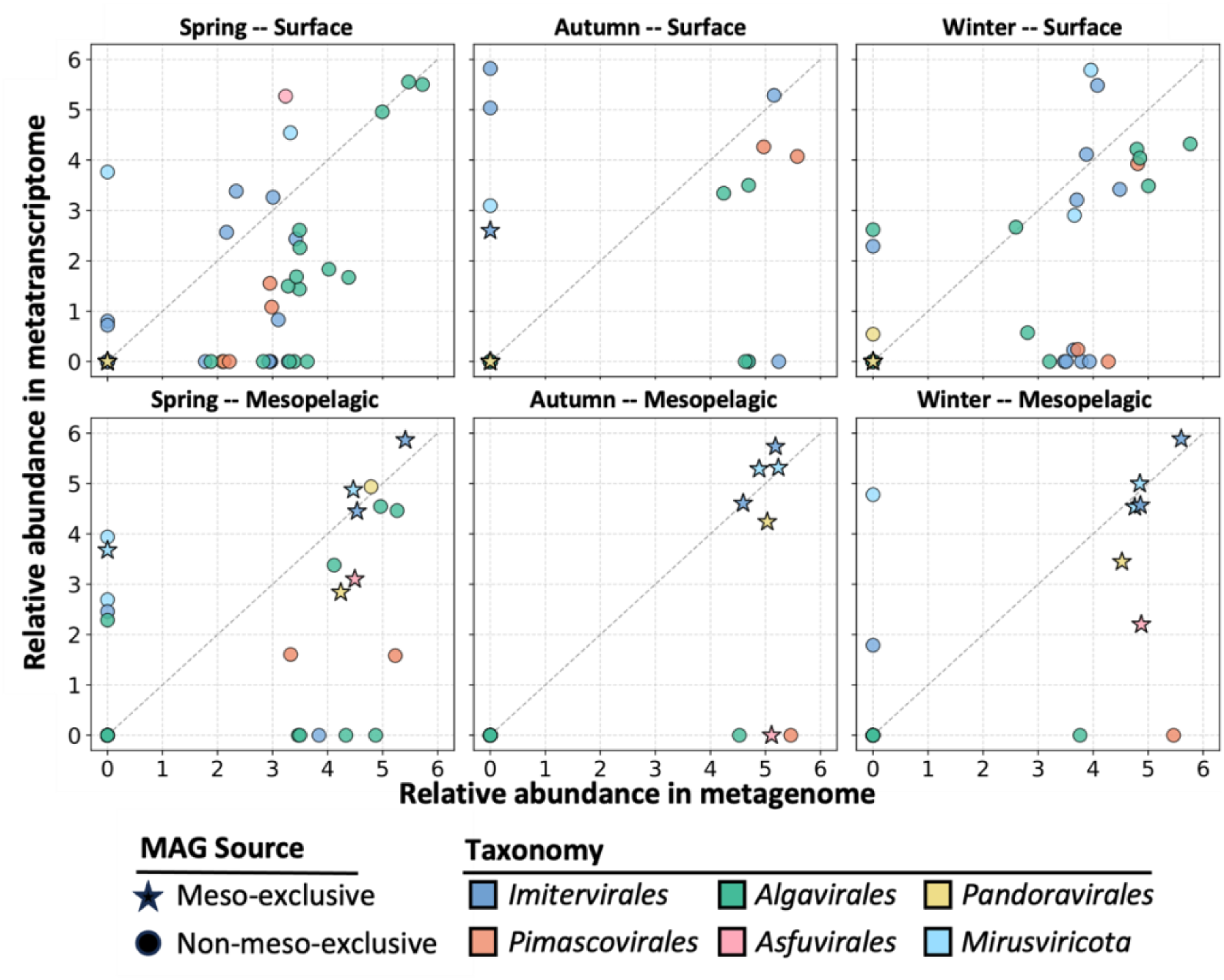
Comparison between metagenomic and metatranscriptomic abundances of Muroto GVMAGs across seasons and depths. Scatterplots showing the relationship between average abundance in metagenomic (X-axis, logTPM+1) and metatranscriptomic (Y-axis, logTPM+1) datasets for Muroto GVMAGs. The first column represents samples from spring, the second from autumn, and the third from winter. The top row corresponds to surface samples, while the bottom row corresponds to mesopelagic samples. Point color indicates taxonomic classification, and shape denotes MAG source: stars represent deep-sea-derived GVMAGs, while circles indicate surface-derived GVMAGs. The dashed diagonal line represents a 1:1 ratio for reference. Surf-derived: GVMAGs assembled from the surface layer. Meso-derived: GVMAGs assembled from the mesopelagic layer.

**Figure S10.**
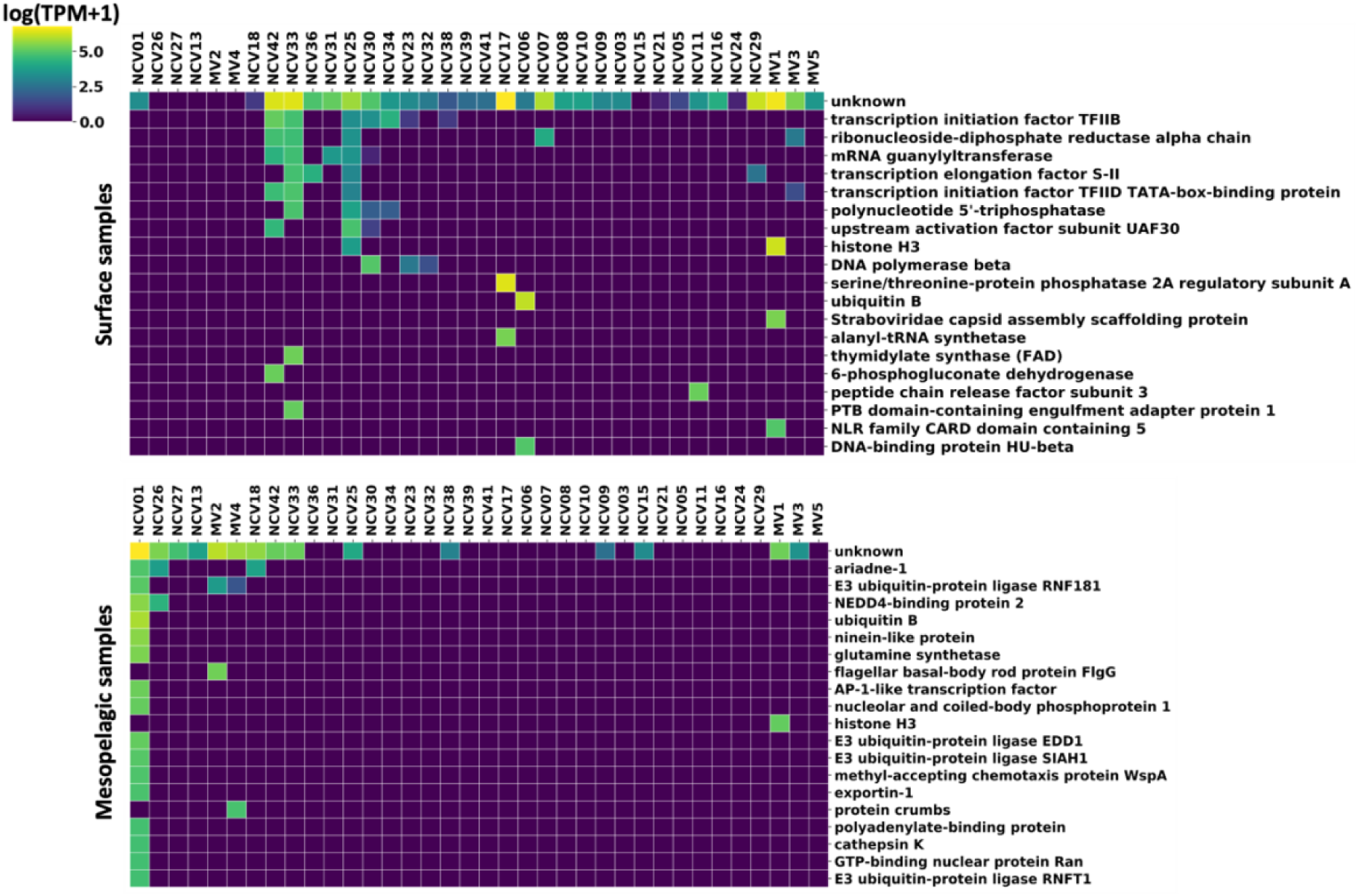
Top 20 KO terms by mean relative abundance among the metatranscriptomic samples in the Muroto GVMAGs. 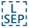 Heatmaps showing the mean relative abundance (logTPM+1) of the top 20 most expressed KEGG orthology (KO) terms across 48 Muroto GVMAGs in (a) surface and (b) mesopelagic metatranscriptomic samples. Each column represents a GVMAG, while each row corresponds to a KO term.

**Figure S11.**
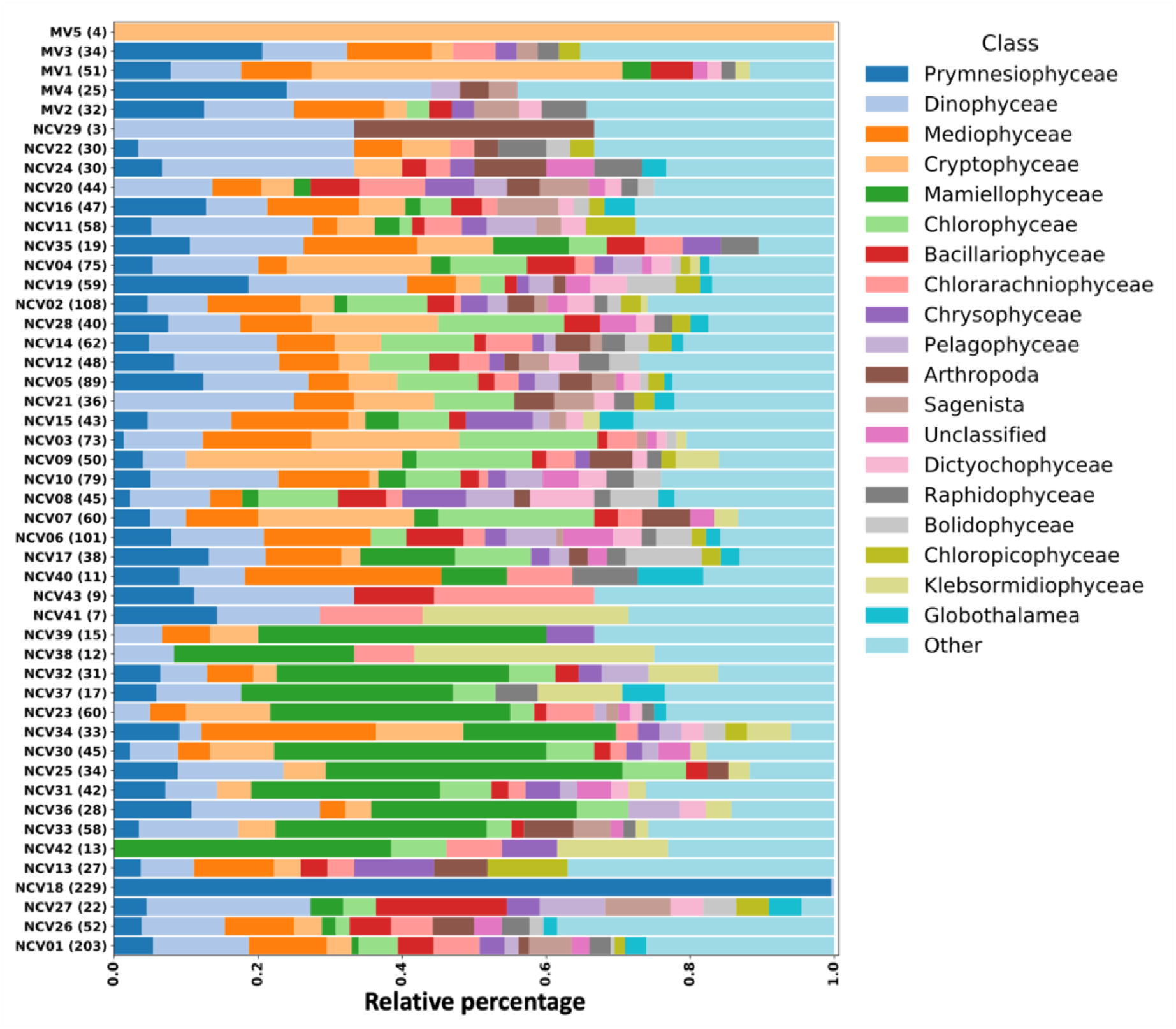
Assignment of eukaryotic-like genes in the Muroto GVMAGs. Each row represents a Muroto GVMAG. Numbers in parentheses following a GVMAG name indicate the total number of genes with hits to eukaryotic taxa in the Marferret database. The X-axis shows the relative percentage of gene hits to corresponding eukaryotic classes. Colors indicating different taxonomic classes. GVMAGs enclosed by black boxes indicate mesopelagic-exclusive GVMAGs.

**Figure S12.**
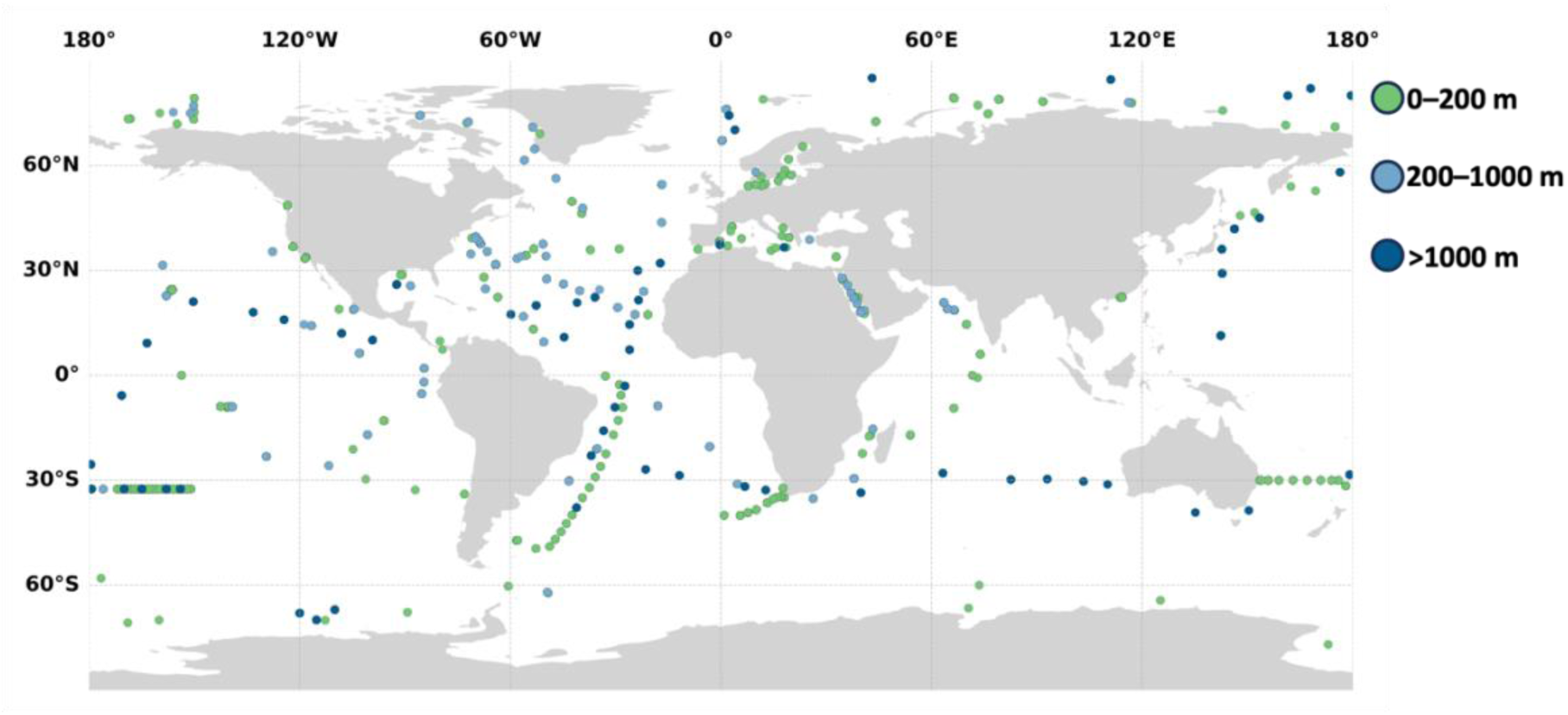
Geographic distribution of 1,890 seawater samples used in this study. Different colored dots represent samples from different depth categories.

**Figure S13.**
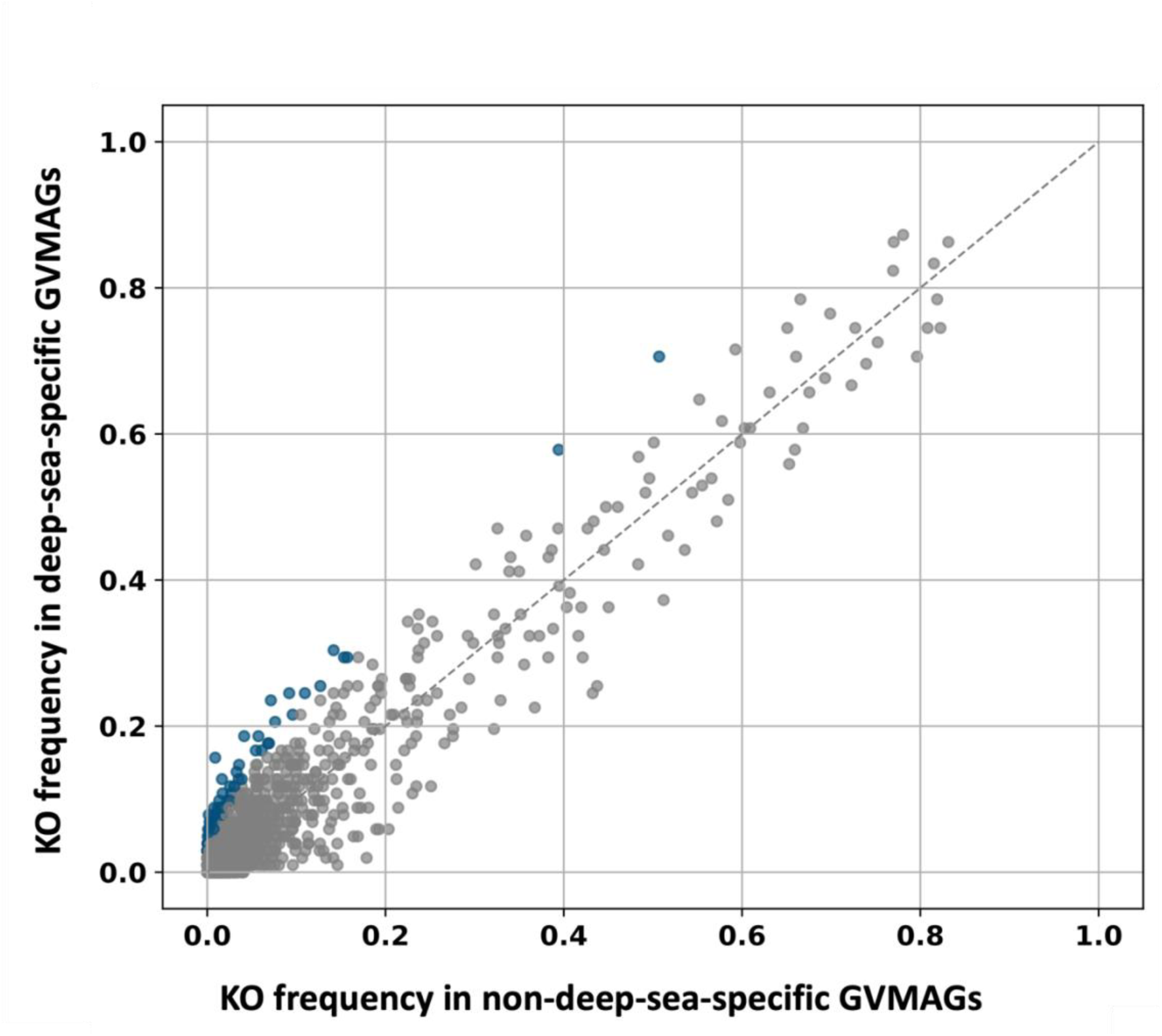
Functional enrichment of deep-sea-specific GVMAGs. Scatterplot showing the comparison of mean KO frequencies between deep-sea-specific and other GVMAGs. Each dot represents a KO term. Those enriched in deep-sea-specific GVMAGs are highlighted in blue. The dashed line indicates a 1:1 ratio reference.

**Figure S14.**
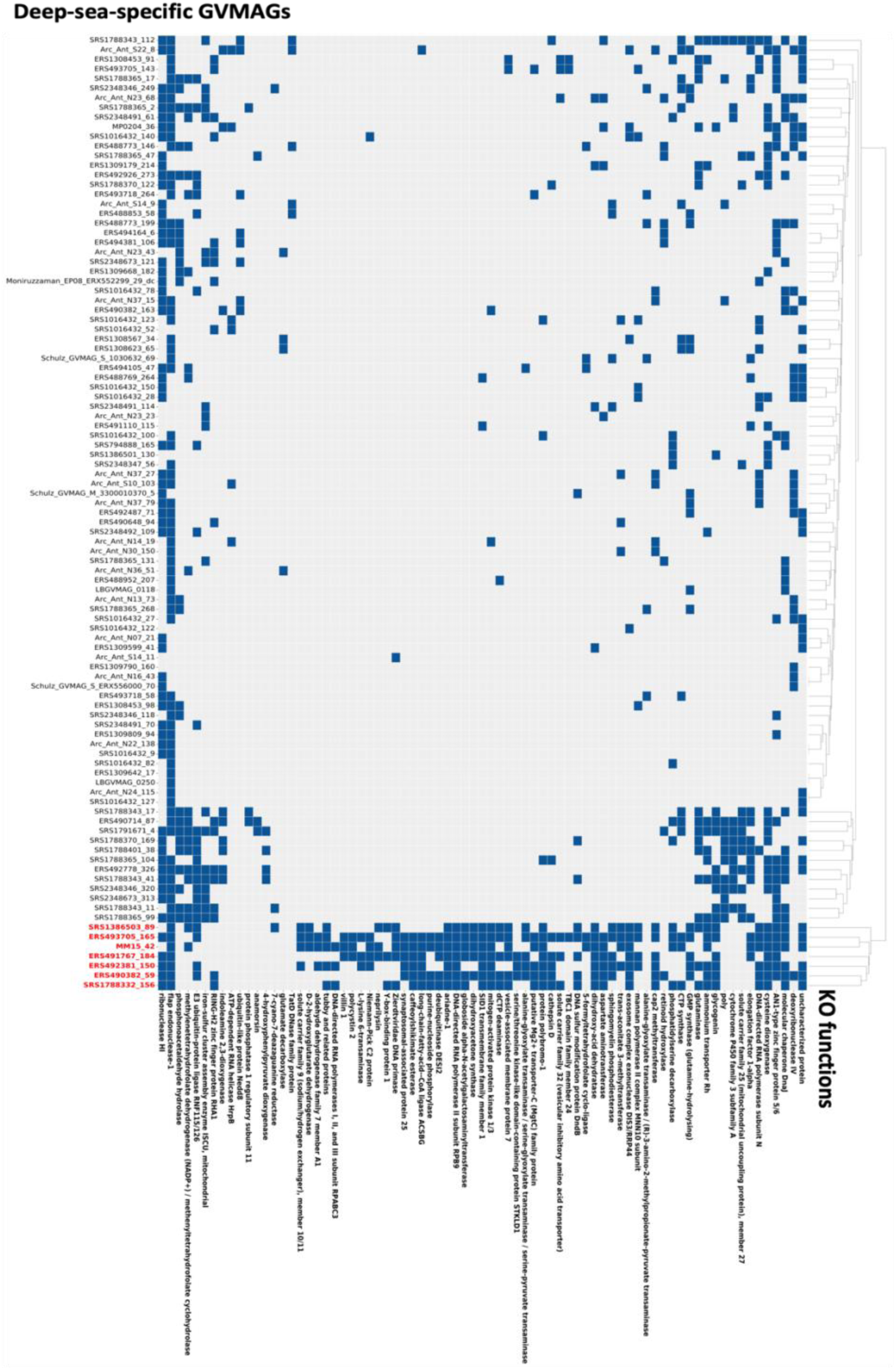
Distribution of deep-sea-specific KO functions across deep-sea-specific GVMAGs. Heatmap showing the presence (blue) or absence (gray) of KO terms enriched in deep-sea-specific GVMAGs. Columns represent KO terms, and rows represent deep-sea-specific GVMAGs. MAG names shown in red indicate GVMAGs in the deep-sea-specific GVMAGs clade described in the main text.

## REFERENCES

1. Schulz F, Abergel C, Woyke T. 2022. Giant virus biology and diversity in the era of genome-resolved metagenomics. Nat Rev Microbiol 20:721–736.

2. Gaïa M, Meng L, Pelletier E, Forterre P, Vanni C, Fernandez-Guerra A, Jaillon O, Wincker P, Ogata H, Krupovic M, Delmont TO. 2023. Mirusviruses link herpesviruses to giant viruses. Nature 2023 616:7958 616:783–789.

3. Archibald J, Chung D, Brask N, Matar S, Gallot-Lavallée L, Pringle E, Duguay B, Blais C, Latimer J, Slamovits C, Leyland B, Rest J, Collier J, McCormick C. 2025. Persistent mirusvirus infection in the marine protist Aurantiochytrium 10.21203/RS.3.RS-5686297/V1.

4. Sun TW, Yang CL, Kao TT, Wang TH, Lai MW, Ku C. 2020. Host range and coding potential of eukaryotic giant viruses. Viruses 12:1–20.

5. Meng L, Endo H, Blanc-Mathieu R, Chaffron S, Hernández-Velázquez R, Kaneko H, Ogata H. 2021. Quantitative Assessment of Nucleocytoplasmic Large DNA Virus and Host Interactions Predicted by Co-occurrence Analyses. mSphere 6.

6. Zhao H, Meng L, Hikida H, Ogata H. 2024. Eukaryotic genomic data uncover an extensive host range of mirusviruses. Curr Biol 34:2633–2643.e3.

7. Wang J, Li L, Lin S, Wang J, Li L, Lin S. 2023. Active viral infection during blooms of a dinoflagellate indicates dinoflagellate-viral co-adaptation.

8. Hevroni G, Vincent F, Ku C, Sheyn U, Vardi A. 2023. Daily turnover of active giant virus infection during algal blooms revealed by single-cell transcriptomics. Sci Adv 9.

9. Funaoka Y, Hiromoto H, Morimoto D, Takahashi M, Wada K, Nagasaki K. 2023. Diversity in Infection Specificity between the Bloom-forming Microalga Heterosigma akashiwo and Its dsDNA Virus, Heterosigma akashiwo Virus. Microbes Environ 38.

10. Rosenwasser S, Ziv C, Creveld SG van, Vardi A. 2016. Virocell Metabolism: Metabolic Innovations During Host-Virus Interactions in the Ocean. Trends Microbiol 24:821–832.

11. Monier A, Pagarete A, De Vargas C, Allen MJ, Read B, Claverie JM, Ogata H. 2009. Horizontal gene transfer of an entire metabolic pathway between a eukaryotic alga and its DNA virus. Genome Res 19:1441–1449.

12. Kaneko H, Blanc-Mathieu R, Endo H, Chaffron S, Delmont TO, Gaia M, Henry N, Hernández-Velázquez R, Nguyen CH, Mamitsuka H, Forterre P, Jaillon O, de Vargas C, Sullivan MB, Suttle CA, Guidi L, Ogata H. 2020. Eukaryotic virus composition can predict the efficiency of carbon export in the global ocean. iScience 24.

13. Hingamp P, Grimsley N, Acinas SG, Clerissi C, Subirana L, Poulain J, Ferrera I, Sarmento H, Villar E, Lima-Mendez G, Faust K, Sunagawa S, Claverie JM, Moreau H, Desdevises Y, Bork P, Raes J, De Vargas C, Karsenti E, Kandels-Lewis S, Jaillon O, Not F, Pesant S, Wincker P, Ogata H. 2013. Exploring nucleo-cytoplasmic large DNA viruses in Tara Oceans microbial metagenomes. ISME J 7:1678–1695.

14. Ha AD, Moniruzzaman M, Aylward FO. 2021. High Transcriptional Activity and Diverse Functional Repertoires of Hundreds of Giant Viruses in a Coastal Marine System. mSystems 6.

15. Carradec Q, Pelletier E, Da Silva C, Alberti A, Seeleuthner Y, Blanc-Mathieu R, Lima-Mendez G, Rocha F, Tirichine L, Labadie K, Kirilovsky A, Bertrand A, Engelen S, Madoui MA, Méheust R, Poulain J, Romac S, Richter DJ, Yoshikawa G, Dimier C, Kandels-Lewis S, Picheral M, Searson S, Acinas SG, Boss E, Follows M, Gorsky G, Grimsley N, Karp-Boss L, Krzic U, Pesant S, Reynaud EG, Sardet C, Sieracki M, Speich S, Stemmann L, Velayoudon D, Weissenbach J, Jaillon O, Aury JM, Karsenti E, Sullivan MB, Sunagawa S, Bork P, Not F, Hingamp P, Raes J, Guidi L, Ogata H, De Vargas C, Iudicone D, Bowler C, Wincker P. 2018. A global ocean atlas of eukaryotic genes. Nat Commun 9.

16. Li Y, Hingamp P, Watai H, Endo H, Yoshida T, Ogata H. 2018. Degenerate PCR Primers to Reveal the Diversity of Giant Viruses in Coastal Waters. Viruses 10.

17. Mihara T, Koyano H, Hingamp P, Grimsley N, Goto S, Ogata H. 2018. Taxon Richness of “Megaviridae” Exceeds those of Bacteria and Archaea in the Ocean. Microbes Environ 33:162–171.

18. Schulz F, Roux S, Paez-Espino D, Jungbluth S, Walsh DA, Denef VJ, McMahon KD, Konstantinidis KT, Eloe-Fadrosh EA, Kyrpides NC, Woyke T. 2020. Giant virus diversity and host interactions through global metagenomics. Nature 2020 578:7795 578:432–436.

19. Endo H, Blanc-Mathieu R, Li Y, Salazar G, Henry N, Labadie K, de Vargas C, Sullivan MB, Bowler C, Wincker P, Karp-Boss L, Sunagawa S, Ogata H. 2020. Biogeography of marine giant viruses reveals their interplay with eukaryotes and ecological functions. Nat Ecol Evol 4:1639–1649.

20. Meng L, Delmont TO, Gaïa M, Pelletier E, Fernàndez-Guerra A, Chaffron S, Neches RY, Wu J, Kaneko H, Endo H, Ogata H. 2023. Genomic adaptation of giant viruses in polar oceans. Nat Commun 14.

21. Bäckström D, Yutin N, Jørgensen SL, Dharamshi J, Homa F, Zaremba-Niedwiedzka K, Spang A, Wolf YI, Koonin E V., Ettema TJG. 2019. Virus genomes from deep sea sediments expand the ocean megavirome and support independent origins of viral gigantism. mBio 10.1128/mBio.02497-18.

22. Prodinger F, Endo H, Takano Y, Li Y, Tominaga K, Isozaki T, Blanc-Mathieu R, Gotoh Y, Hayashi T, Taniguchi E, Nagasaki K, Yoshida T, Ogata H. 2022. Year-round dynamics of amplicon sequence variant communities differ among eukaryotes, Imitervirales and prokaryotes in a coastal ecosystem. FEMS Microbiol Ecol 97.

23. Fang Y, Meng L, Xia J, Gotoh Y, Hayashi T, Nagasaki K, Endo H, Okazaki Y, Ogata H. 2025. Genome-resolved year-round dynamics reveal a broad range of giant virus microdiversity. mSystems 10.

24. Fromm A, Hevroni G, Vincent F, Schatz D, Martinez-Gutierrez CA, Aylward FO, Vardi A. 2024. Single-cell RNA-seq of the rare virosphere reveals the native hosts of giant viruses in the marine environment. Nat Microbiol 9:1619–1629.

25. Xia J, Kameyama S, Prodinger F, Yoshida T, Cho KH, Jung J, Kang SH, Yang EJ, Ogata H, Endo H. 2022. Tight association between microbial eukaryote and giant virus communities in the Arctic Ocean. Limnol Oceanogr 67:1343–1356.

26. Abrahão J, Silva L, Silva LS, Khalil JYB, Rodrigues R, Arantes T, Assis F, Boratto P, Andrade M, Kroon EG, Ribeiro B, Bergier I, Seligmann H, Ghigo E, Colson P, Levasseur A, Kroemer G, Raoult D, La Scola B. 2018. Tailed giant Tupanvirus possesses the most complete translational apparatus of the known virosphere. Nat Commun 9.

27. Mizuno CM, Ghai R, Saghaï A, López-García P, Rodriguez-Valeraa F. 2016. Genomes of Abundant and Widespread Viruses from the Deep Ocean. mBio 7.

28. Jian H, Yi Y, Wang J, Hao Y, Zhang M, Wang S, Meng C, Zhang Y, Jing H, Wang Y, Xiao X. 2021. Diversity and distribution of viruses inhabiting the deepest ocean on Earth. ISME J 15:3094–3110.

29. Sheam MM, Luo E. 2025. Vertical transport and spatiotemporal dynamics of giant viruses in the North Pacific Subtropical Gyre. ISME J 10.1093/ISMEJO/WRAF094.

30. Acinas SG, Sánchez P, Salazar G, Cornejo-Castillo FM, Sebastián M, Logares R, Royo-Llonch M, Paoli L, Sunagawa S, Hingamp P, Ogata H, Lima-Mendez G, Roux S, González JM, Arrieta JM, Alam IS, Kamau A, Bowler C, Raes J, Pesant S, Bork P, Agustí S, Gojobori T, Vaqué D, Sullivan MB, Pedrós-Alió C, Massana R, Duarte CM, Gasol JM. 2021. Deep ocean metagenomes provide insight into the metabolic architecture of bathypelagic microbial communities. Commun Biol 4.

31. Ha AD, Moniruzzaman M, Aylward FO. 2023. Assessing the biogeography of marine giant viruses in four oceanic transects. ISME 2023.01.30.526306.

32. Zhang L, Meng L, Fang Y, Ogata H, Okazaki Y. 2024. Spatiotemporal dynamics of giant viruses within a deep freshwater lake reveal a distinct dark-water community. ISME J 18:182.

33. Wu J, Gao W, Zhang W, Meldrum DR. 2011. Optimization of whole-transcriptome amplification from low cell density deep-sea microbial samples for metatranscriptomic analysis. J Microbiol Methods 84:88–93.

34. Steiner PA, De Corte D, Geijo J, Mena C, Yokokawa T, Rattei T, Herndl GJ, Sintes E. 2019. Highly variable mRNA half-life time within marine bacterial taxa and functional genes. Environ Microbiol 21:3873–3884.

35. Guo Y.J. 1991. The Kuroshio. Part II. Primary productivity and phytoplankton. Oceanogr Mar Biol Annu Rev.

36. Yamaguchi T, Inoue T, Hirakawa M, Abe S, Ishii K, Kagoura T, Fujiwara M. 2003. Deep-sea water suction technology. Furukawa Review 75–80.

37. Kang Y, Kawamoto J, Kaneda S, Aritome K, Sakurai K. 2004. Characterization of Selenium in the Deep Ocean Water Pumped up at Muroto, Japan. https://www.researchgate.net/publication/306472302_Characterization_of_Selenium_in_the_Deep_Ocean_Water_Pumped_up_at_Muroto_Japan. Retrieved 27 March 2025.

38. Nishimura Y, Yoshizawa S. 2022. The OceanDNA MAG catalog contains over 50,000 prokaryotic genomes originated from various marine environments. Scientific Data 2022 9:1 9:1–11.

39. Collier JL, Rest JS, Gallot-Lavallée L, Lavington E, Kuo A, Jenkins J, Plott C, Pangilinan J, Daum C, Grigoriev I V., Filloramo G V., Novák Vanclová AMG, Archibald JM. 2023. The protist Aurantiochytrium has universal subtelomeric rDNAs and is a host for mirusviruses. Curr Biol 33:5199–5207.e4.

40. Prokopowich CD, Gregory TR, Crease TJ. 2003. The correlation between rDNA copy number and genome size in eukaryotes. Genome 46:48–50.

41. Zhu F, Massana R, Not F, Marie D, Vaulot D. 2005. Mapping of picoeucaryotes in marine ecosystems with quantitative PCR of the 18S rRNA gene. FEMS Microbiol Ecol 52:79–92.

42. Giner CR, Pernice MC, Balagué V, Duarte CM, Gasol JM, Logares R, Massana R. 2020. Marked changes in diversity and relative activity of picoeukaryotes with depth in the world ocean. ISME J 14:437–449.

43. Ogata H, Toyoda K, Tomaru Y, Nakayama N, Shirai Y, Claverie JM, Nagasaki K. 2009. Remarkable sequence similarity between the dinoflagellate-infecting marine girus and the terrestrial pathogen African swine fever virus. Virol J 6.

44. Da Cunha V, Gaia M, Ogata H, Jaillon O, Delmont TO, Forterre P. 2022. Giant Viruses Encode Actin-Related Proteins. Mol Biol Evol 39.

45. Kijima S, Hikida H, Delmont TO, Gaïa M, Ogata H. 2024. Complex Genomes of Early Nucleocytoviruses Revealed by Ancient Origins of Viral Aminoacyl-tRNA Synthetases. Mol Biol Evol 41.

46. Wu B, Fan T, Chen X, He Y, Wang H. 2024. The class III phosphatidylinositol 3-kinase VPS34 supports EV71 replication by promoting viral replication organelle formation. J Virol 98.

47. Tang Q, Wu P, Chen H, Li G. 2018. Pleiotropic roles of the ubiquitin-proteasome system during viral propagation. Life Sci 207:350.

48. Buscaglia M, Iriarte JL, Schulz F, Díez B. 2024. Adaptation strategies of giant viruses to low-temperature marine ecosystems. ISME J 18:162.

49. Lamb DC, Follmer AH, Goldstone J V., Nelson DR, Warrilow AG, Price CL, True MY, Kelly SL, Poulos TL, Stegeman JJ. 2019. On the occurrence of cytochrome P450 in viruses. Proc Natl Acad Sci U S A 116:12343–12352.

50. Lamb DC, Goldstone J V., Belhaouari DB, Andréani J, Farooqi A, Allen MJ, Kelly SL, La Scola B, Stegeman JJ. 2025. Cytochrome b5 occurrence in giant and other viruses belonging to the phylum Nucleocytoviricota. npj Viruses 2025 3:1 3:1–11.

51. Hwang DM, Dempsey A, Tan KT, Liew CC. 1996. A modular domain of NiFu, a nitrogen fixation cluster protein, is highly conserved in evolution. J Mol Evol 43:536–540.

52. Moniruzzaman M, Martinez-Gutierrez CA, Weinheimer AR, Aylward FO. 2020. Dynamic genome evolution and complex virocell metabolism of globally-distributed giant viruses. Nat Commun 11:1–11.

53. Kim SK, Makino K, Amemura M, Shinagawa H, Nakata A. 1993. Molecular analysis of the phoH gene, belonging to the phosphate regulon in Escherichia coli. J Bacteriol 175:1316–1324.

54. Okazaki Y, Nguyen TT, Nishihara A, Endo H, Ogata H, Nakano SI, Tamaki H. 2023. A Fast and Easy Method to Co-extract DNA and RNA from an Environmental Microbial Sample. Microbes Environ 38.

55. Comeau AM, Li WKW, Tremblay JÉ, Carmack EC, Lovejoy C. 2011. Arctic Ocean microbial community structure before and after the 2007 record sea ice minimum. PLoS One 6.

56. Bolyen E, Rideout JR, Dillon MR, Bokulich NA, Abnet CC, Al-Ghalith GA, Alexander H, Alm EJ, Arumugam M, Asnicar F, Bai Y, Bisanz JE, Bittinger K, Brejnrod A, Brislawn CJ, Brown CT, Callahan BJ, Caraballo-Rodríguez AM, Chase J, Cope EK, Da Silva R, Diener C, Dorrestein PC, Douglas GM, Durall DM, Duvallet C, Edwardson CF, Ernst M, Estaki M, Fouquier J, Gauglitz JM, Gibbons SM, Gibson DL, Gonzalez A, Gorlick K, Guo J, Hillmann B, Holmes S, Holste H, Huttenhower C, Huttley GA, Janssen S, Jarmusch AK, Jiang L, Kaehler BD, Kang K Bin, Keefe CR, Keim P, Kelley ST, Knights D, Koester I, Kosciolek T, Kreps J, Langille MGI, Lee J, Ley R, Liu YX, Loftfield E, Lozupone C, Maher M, Marotz C, Martin BD, McDonald D, McIver LJ, Melnik A V., Metcalf JL, Morgan SC, Morton JT, Naimey AT, Navas-Molina JA, Nothias LF, Orchanian SB, Pearson T, Peoples SL, Petras D, Preuss ML, Pruesse E, Rasmussen LB, Rivers A, Robeson MS, Rosenthal P, Segata N, Shaffer M, Shiffer A, Sinha R, Song SJ, Spear JR, Swafford AD, Thompson LR, Torres PJ, Trinh P, Tripathi A, Turnbaugh PJ, Ul-Hasan S, van der Hooft JJJ, Vargas F, Vázquez-Baeza Y, Vogtmann E, von Hippel M, Walters W, Wan Y, Wang M, Warren J, Weber KC, Williamson CHD, Willis AD, Xu ZZ, Zaneveld JR, Zhang Y, Zhu Q, Knight R, Caporaso JG. 2019. Reproducible, interactive, scalable and extensible microbiome data science using QIIME 2. Nature Biotechnology 2019 37:8 37:852–857.

57. Callahan BJ, McMurdie PJ, Rosen MJ, Han AW, Johnson AJA, Holmes SP. 2016. DADA2: High-resolution sample inference from Illumina amplicon data. Nat Methods 13:581–583.

58. Bokulich NA, Kaehler BD, Rideout JR, Dillon M, Bolyen E, Knight R, Huttley GA, Gregory Caporaso J. 2018. Optimizing taxonomic classification of marker-gene amplicon sequences with QIIME 2’s q2-feature-classifier plugin. Microbiome 6:1–17.

59. R Core Team. 2024. _R: a language and environment for statistical computing_.

60. Chen S, Zhou Y, Chen Y, Gu J. 2018. fastp: an ultra-fast all-in-one FASTQ preprocessor. Bioinformatics 34:i884–i890.

61. Li D, Liu CM, Luo R, Sadakane K, Lam TW. 2015. MEGAHIT: an ultra-fast single-node solution for large and complex metagenomics assembly via succinct de Bruijn graph. Bioinformatics 31:1674–1676.

62. Langmead B, Wilks C, Antonescu V, Charles R. 2019. Scaling read aligners to hundreds of threads on general-purpose processors. Bioinformatics 35:421–432.

63. Kang DD, Li F, Kirton E, Thomas A, Egan R, An H, Wang Z. 2019. MetaBAT 2: An adaptive binning algorithm for robust and efficient genome reconstruction from metagenome assemblies. PeerJ 2019:e7359.

64. Hyatt D, Chen GL, LoCascio PF, Land ML, Larimer FW, Hauser LJ. 2010. Prodigal: Prokaryotic gene recognition and translation initiation site identification. BMC Bioinformatics 11:1–11.

65. Camargo AP, Roux S, Schulz F, Babinski M, Xu Y, Hu B, Chain PSG, Nayfach S, Kyrpides NC. 2023. Identification of mobile genetic elements with geNomad. Nature Biotechnology 2023 42:8 42:1303–1312.

66. Yutin N, Wolf YI, Raoult D, Koonin E V. 2009. Eukaryotic large nucleo-cytoplasmic DNA viruses: clusters of orthologous genes and reconstruction of viral genome evolution. Virol J 6.

67. Aylward FO, Moniruzzaman M. 2021. ViralRecall-A Flexible Command-Line Tool for the Detection of Giant Virus Signatures in ’Omic Data. Viruses 13.

68. Guo J, Bolduc B, Zayed AA, Varsani A, Dominguez-Huerta G, Delmont TO, Pratama AA, Gazitúa MC, Vik D, Sullivan MB, Roux S. 2021. VirSorter2: a multi-classifier, expert-guided approach to detect diverse DNA and RNA viruses. Microbiome 9:1–13.

69. Von Meijenfeldt FAB, Arkhipova K, Cambuy DD, Coutinho FH, Dutilh BE. 2019. Robust taxonomic classification of uncharted microbial sequences and bins with CAT and BAT. Genome Biol 20:1–14.

70. Eren AM, Esen OC, Quince C, Vineis JH, Morrison HG, Sogin ML, Delmont TO. 2015. Anvi’o: an advanced analysis and visualization platform for ’omics data. PeerJ 3.

71. Mistry J, Finn RD, Eddy SR, Bateman A, Punta M. 2013. Challenges in homology search: HMMER3 and convergent evolution of coiled-coil regions. Nucleic Acids Res 41.

72. Nayfach S, Camargo AP, Schulz F, Eloe-Fadrosh E, Roux S, Kyrpides NC. 2020. CheckV assesses the quality and completeness of metagenome-assembled viral genomes. Nature Biotechnology 2020 39:5 39:578–585.

73. Olm MR, Brown CT, Brooks B, Banfield JF. 2017. dRep: a tool for fast and accurate genomic comparisons that enables improved genome recovery from metagenomes through de-replication. The ISME Journal 2017 11:12 11:2864–2868.

74. Mortazavi A, Williams BA, McCue K, Schaeffer L, Wold B. 2008. Mapping and quantifying mammalian transcriptomes by RNA-Seq. Nat Methods 5:621–628.

75. Aroney STN, Newell RJP, Nissen JN, Camargo AP, Tyson GW, Woodcroft BJ. 2025. CoverM: read alignment statistics for metagenomics. Bioinformatics 41.

76. Kopylova E, Noé L, Touzet H. 2012. SortMeRNA: fast and accurate filtering of ribosomal RNAs in metatranscriptomic data. Bioinformatics 28:3211–3217.

77. Patro R, Duggal G, Love MI, Irizarry RA, Kingsford C. 2017. Salmon provides fast and bias-aware quantification of transcript expression. Nature Methods 2017 14:4 14:417– 419.

78. Moniruzzaman M, Martinez-Gutierrez CA, Weinheimer AR, Aylward FO. 2020. Dynamic genome evolution and complex virocell metabolism of globally-distributed giant viruses. Nature Communications 2020 11:1 11:1–11.

79. Sievers F, Higgins DG. 2014. Clustal Omega, accurate alignment of very large numbers of sequences. Methods Mol Biol 1079:105–116.

80. Capella-Gutiérrez S, Silla-Martínez JM, Gabaldón T. 2009. trimAl: a tool for automated alignment trimming in large-scale phylogenetic analyses. Bioinformatics 25:1972–1973.

81. Price MN, Dehal PS, Arkin AP. 2010. FastTree 2 – Approximately Maximum-Likelihood Trees for Large Alignments. PLoS One 5:e9490.

82. Aylward FO, Moniruzzaman M, Ha AD, Koonin E V. 2021. A phylogenomic framework for charting the diversity and evolution of giant viruses. PLoS Biol 19:e3001430.

83. Ha AD, Aylward FO. 2024. Automated classification of giant virus genomes using a random forest model built on trademark protein families. npj Viruses 2024 2:1 2:1–9.

84. Benjamini Y, Hochberg Y. 1995. Controlling the False Discovery Rate: A Practical and Powerful Approach to Multiple Testing. Journal of the Royal Statistical Society: Series B (Methodological) 57:289–300.

85. Aramaki T, Blanc-Mathieu R, Endo H, Ohkubo K, Kanehisa M, Goto S, Ogata H. 2020. KofamKOALA: KEGG Ortholog assignment based on profile HMM and adaptive score threshold. Bioinformatics 36:2251–2252.

86. Mistry J, Chuguransky S, Williams L, Qureshi M, Salazar GA, Sonnhammer ELL, Tosatto SCE, Paladin L, Raj S, Richardson LJ, Finn RD, Bateman A. 2021. Pfam: The protein families database in 2021. Nucleic Acids Res 49:D412–D419.

87. Buchfink B, Reuter K, Drost HG. 2021. Sensitive protein alignments at tree-of-life scale using DIAMOND. Nature Methods 2021 18:4 18:366–368.

88. Groussman RD, Blaskowski S, Coesel SN, Armbrust E V. 2023. MarFERReT, an open-source, version-controlled reference library of marine microbial eukaryote functional genes. Scientific Data 2023 10:1 10:1–19.

89. Kazlauskas D, Krupovic M, Guglielmini J, Forterre P, Venclovas CS. 2020. Diversity and evolution of B-family DNA polymerases. Nucleic Acids Res 48:10142–10156.

90. Nguyen LT, Schmidt HA, Von Haeseler A, Minh BQ. 2015. IQ-TREE: A Fast and Effective Stochastic Algorithm for Estimating Maximum-Likelihood Phylogenies. Mol Biol Evol 32:268–274.

91. Hoang DT, Chernomor O, Von Haeseler A, Minh BQ, Vinh LS. 2018. UFBoot2: Improving the Ultrafast Bootstrap Approximation. Mol Biol Evol 35:518–522.

92. Letunic I, Bork P. 2024. Interactive Tree of Life (iTOL) v6: recent updates to the phylogenetic tree display and annotation tool. Nucleic Acids Res 52:W78–W82.

